# Celldetective: an AI-enhanced image analysis tool for unraveling dynamic cell interactions

**DOI:** 10.1101/2024.03.15.585250

**Authors:** Rémy Torro, Beatriz Díaz-Bello, Dalia El Arawi, Ksenija Dervanova, Lorna Ammer, Florian Dupuy, Patrick Chames, Kheya Sengupta, Laurent Limozin

## Abstract

Analysis of multimodal and multidimensional data capturing dynamic interactions between diverse cell populations is a current challenge in bioimaging, especially in the context of immunology and immunotherapy research. Here, we introduce Celldetective, an open-source Python-based software tool designed for high-performance, end-to-end analysis of image-based *in vitro* immune and immunotherapy assays. Celldetective is purpose-built for multicondition, 2D multi-channel time-lapse microscopy of mixed cell populations. Although it is optimised for the needs of immunology assays, it is nevertheless broadly applicable to any biological system involving interacting cell populations. The software seamlessly integrates AI-based segmentation, tracking, and automated single-cell event detection, all within an intuitive graphical interface that supports interactive visualisation, annotation, and training options. We showcase its capabilities with original datasets of single immune effector cell interactions with an activating surface mediated by bispecific antibodies, and pairwise interactions in antibody-dependent cell cytotoxicity events.

## Introduction

An overarching goal of post-genomic biomedical research is to decipher cell states and transitions as well as cell interactions in functional dynamic assays [1, 2]. This requires high-throughput and high-resolution experimental methods supported by advanced and powerful analyses [3, 4]. The field of immunology presents specific challenges, related to complex cell states characterised by transient and heterogeneous receptor expression, highly motile behaviour and immune function relying on dynamic cell-cell interactions [5, 6]. These features are largely out of reach in ensemble biochemical measurements [7] as well as low time-resolution single-cell multi-omic data [8].

Microscopy-based *in vitro* assays can bridge temporal and spatial scales, helping to decipher lymphocyte activation from molecular to cellular scales [9–11]. Combining high-content and high-spatial-resolution, imaging flow cytometry was applied to reconstruct disease progression on single immune-cell populations [12] or to functionally characterise therapeutic antibodies against liquid tumours [13]. The analysis is however limited to single cells or cell doublets. Multichannel fluorescence microscopy was successfully applied to image-based profiling of human T lymphocytes and Natural Killer cells (NK) [14] or therapeutic targets in drug discovery [15], but it provided limited dynamic information on the system. In some cases, nanowell grids were used to organise cell contacts and facilitate image analysis [16, 17] at the expense of imposing less physiological conditions.

High-content, multi-channel spatiotemporal imaging of living cells provides rich, dynamic and multiscale information, from molecular to cell population scale [18]. In single-cell image sequences, time-dependent signals contain functional information, traditionally collected or treated manually in immunological settings [19]. Recent methods performed profiling on time-dependent data for cellular phenotyping, but without measuring single events in time [20]. Functional dynamic information like calcium imaging can also be extracted for immune cell classification [21, 22] or to interpret the brain’s neuronal activity [23]. Most recent works on cell dynamics, especially in the context of cancer immunotherapy, involve specialised microscopy techniques and complex analysis protocols [24–26], sometimes using proprietary software [27, 28].

In recent years, machine learning and deep learning (DL) have revolutionised the field of image analysis [12, 13, 29–33]. A critical step in the context of cell images is segmentation, for which DL tools are highly successful [34–39]. However, they often rely on pretrained models for which the original dataset is not always accessible or can be too remote from users’ data [40], limiting adaptability. For dynamic data, cell tracking is also crucial and can be performed by available tools [41–43]. Recent methods that integrate profiling with time-dependent information significantly enhance cell tracking performance [44–46]. However, current software tools associating the two tasks are mostly applied to the analysis of cell lineage [42, 47–49]. Most popular free bioimage analysis software tools like Fiji [50], Icy [51] or Cellprofiler [52] were not originally designed for DL methods [53], relying on extensions and plugins to bring in these new methods, or were not conceived for time-dependent data [54].

Finally, end-to-end analysis tools accessible to experimentalists, adapted for both images and time series are currently lacking, preventing dataset curation and DL-based event detection. With the exception of MIA [55], existing partial solutions require coding skills [56] or proficiency in integrating/mastering disparate tools [57].

We introduce Celldetective, an open-source software tool designed to tackle the challenge of analysing image-based *in vitro* immune and immunotherapy assays. It addresses the problem of cell interactions by extracting and interpreting single-cell behaviour for user-defined cell populations in a 2D co-culture, with pairwise descriptions linking any two populations. The software adapts to varying imaging conditions and cell types, incorporating dataset curation and deep learning model training features directly within the graphical user interface (GUI). Celldetective organises and automates complicated and time-consuming processes while providing a user-friendly, no-code GUI, making it accessible to experimental scientists without programming expertise.

We tested the software on two applications in immunology and immunotherapy: i) measuring immune cell behaviour in contact with an antigen-presenting surface using a label-free microscopy technique, and ii) assessing dynamic cell-cell interactions in a multiwell co-culture of immune and target cells, an established cytotoxic assay, using multicolour fluorescence microscopy.

## Results

### Software analysis pipeline

Celldetective is implemented as a Python package with a PyQt5-based GUI, available through the PyPi package repository and regularly updated. The source code can be accessed on GitHub (https://github.com/celldetective/celldetective), while all models, datasets, and demos are hosted in a public Zenodo repository (https://zenodo.org/records/13907294). Extensive documentation is available online (https://celldetective.readthedocs.io). The software is designed to run locally on a single computer equipped with either a graphics processing unit (GPU) or multiple processors. Image processing is conducted frame by frame with garbage collection to significantly reduce memory requirements, allowing it to operate on most laboratory computers. Celldetective is compatible with multi-channel time-lapse microscopy image stack files in TIF/OME.TIF format, which are typically generated by acquisition software such as Micro-Manager [58].

As illustrated in Fig. 1, the software input consists of an experimental project that organises raw data into a two-level folder structure emulating a multiwell plate, commonly used in *in vitro* immune assays (startup window in Fig. S1A). At the top level, folders represent wells (or biological conditions), while at the second level, folders represent positions (a field of view associated with the biological condition). Experiment metadata – including channel names and order, spatiotemporal calibration, and biological condition labels – are stored in a configuration file created during project setup (Fig. S1B). This structure organises the experimental data by biological conditions and fields of view, facilitating the processing of large data volumes.

**Fig. 1.**
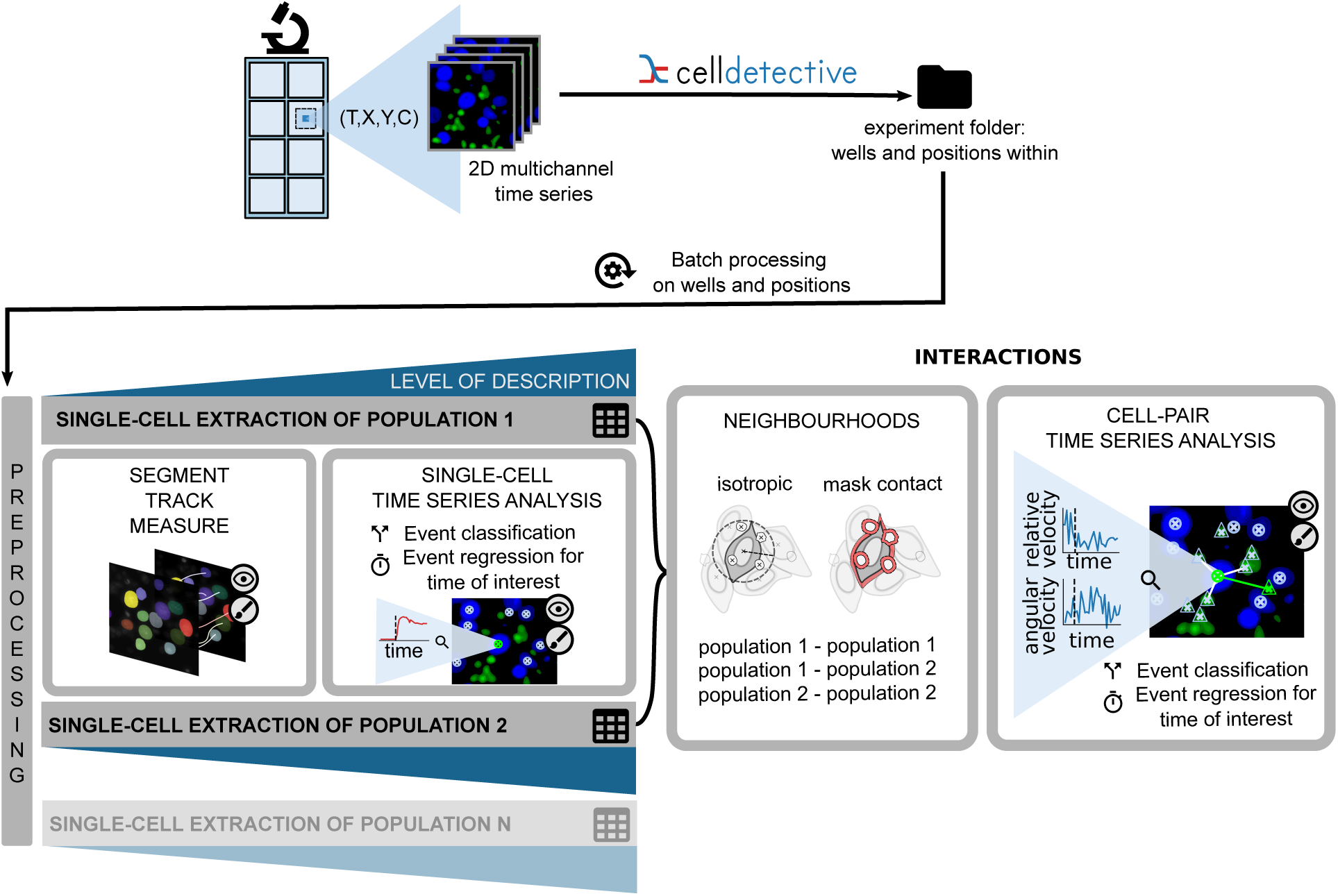
Software Overview: an end-to-end GUI pipeline, from left to right, for studying interactions between cell populations, here illustrated with target and effector populations. After loading an experiment project that mimics a multiwell plate structure, the user can apply preprocessing steps to the 2D time-lapse microscopy images before segmentation. Target and effector cells are then segmented, tracked, and measured independently. Events are detected from the resulting time series, and the co-culture images are distilled into tables of single-cell measurements. The neighbourhood module links cells in spatial proximity, and the cell-pair signal analysis framework facilitates the investigation of interactions between cell pairs. Eye and brush icons indicate steps where visual control and corrections are possible, with an appropriate viewer.

Upon loading a project, a modular interface organises all possible user actions (Fig. S1C, with single-cell extraction modules illustrated in Fig. S2). Image preprocessing enables users to transform images prior to segmentation (Fig. S3). Independent processing blocks drive the single-cell extraction for each user-defined population. Population-specific segmentation, optional tracking and measurements, reduce the image stacks into tables, where each row represents a single cell at a specific time point. With tracking information available, single-cell measurements are structured and interpreted as time series to detect events. Alternatively, users can classify cells at each time point from their measurements. A viewer is integrated at each step for quality control, error correction, or data annotation. Notably, both segmentation masks and cell tracks can be interactively corrected within the same napari environment, enabling a seamless curation workflow that is directly linked back to the analysis pipeline. The final module addresses cell interactions within or across populations, enabling users to describe and annotate cell pairs. Throughout the process, users can plot and compare single-cell or cell pair measurement distributions via a table exploration module designed for manipulating tracks. Additional analysis modules allow users to perform survival studies and represent synchronised time series over cell populations.

Rather than reinventing existing efficient solutions, we incorporated into Celldetective established open-source deep learning segmentation methods StarDist [36] and Cellpose [37] (see model list in Tab. S1), alongside the Bayesian tracker bTrack [42] and the trackpy linking algorithm [59]. The software provides intuitive graphical options for tuning input and control parameters of these algorithms. The diverse software functionalities are listed and compared to existing software in Table 1. Notably, Celldetective stands out by offering training and transfer options in segmentation, enabling detection of a population of interest within a mixed population, as well as time series annotation and survival analysis, not currently integrated in any other software. These features will be illustrated in the two biological applications described below.

**Table 1.** Comparative table of software functionalities with a selection of available solutions. A ✓is attributed if the task can be carried out without coding. The use of an integrated solution or plugin is indicated in parentheses. ^∗^Cell-cycle stages only. ^†^Static per-frame neighbour measurement only; no dynamic pair tracking or time series analysis.

| Software feature | ImageJ/Fiji | CellProfiler | CellACDC | MIA | Celldetective |
| --- | --- | --- | --- | --- | --- |
| Ref. | [50] | [52] | [48] | [55] | this work |
| Traditional segmentation | ✓ | ✓ | ✓ | ✗ | ✓ |
| DL segmentation | ✓ | ✓ | ✓ | ✓ | ✓ |
| Segmentation correction | ✓<br>(Labkit) | ✗ | ✓ | ✗ | ✓<br>(napari) |
| Training annotation | ✓<br>(Labkit) | ✓<br>(GIMP) | ✓ | ✓ | ✓<br>(napari) |
| Training | ✗ | ✗ | ✗ | ✓ | ✓ |
| Transfer | ✗ | ✗ | ✗ | ✓ | ✓ |
| Tracking | ✓<br>(TrackMate) | ✓ | ✓ | ✓ | ✓<br>(bTrack, Trackpy) |
| Track visualisation | ✓<br>(TrackMate) | ✗ | ✓ | ✗ | ✓<br>(napari) |
| Track correction | ✓<br>(TrackMate) | ✗ | ✓ | ✗ | ✓<br>(napari) |
| Classification from measurements | ✗ | ✓ | ✗ | ✗ | ✓ |
| Time series annotations | ✗ | ✗ | ✓* | ✗ | ✓ |
| Interaction analysis | ✗ | ✓† | ✗ | ✗ | ✓ |
| Experiment manager | ✗ | ✗ | ✓ | ✗ | ✓ |
| Multi-condition data exploration | ✗ | ✓<br>(CP Analyst) | ✗ | ✗ | ✓ |
| Survival analysis | ✗ | ✗ | ✗ | ✗ | ✓ |

### High-throughput and time-resolved analysis of cell adhesion and spreading responses

Bispecific antibodies (bsAb) act as molecular bridges to force the contact between immune effector cells and cancer cells. Despite the therapeutic interest of these constructs to improve cancer cell killing, their mode of action is poorly understood. In this first application, we study how a bsAb controls the interaction between Natural Killer (NK) cells and a surrogate cancer cell surface [60, 62]. The bsAb bridges the NK cell’s activating receptor CD16 and the Human Epidermal Growth Factor Receptor

2 (HER2) coated on the surface. Cell-surface contact and spreading is monitored using reflection interference contrast microscopy (RICM), a label-free technique sensitive to submicrometer distances between the cell membrane and the surface [11, 60, 61, 63]. The assay consists in depositing freshly isolated human primary NK cells on the HER2 surface, in the presence of known concentrations of bsAb. RICM imaging results in variations of grey levels under the cell in a grey background and provides a high spatiotemporal signal related to cell-surface contact without labelling (Fig. 2A).

**Fig. 2.**
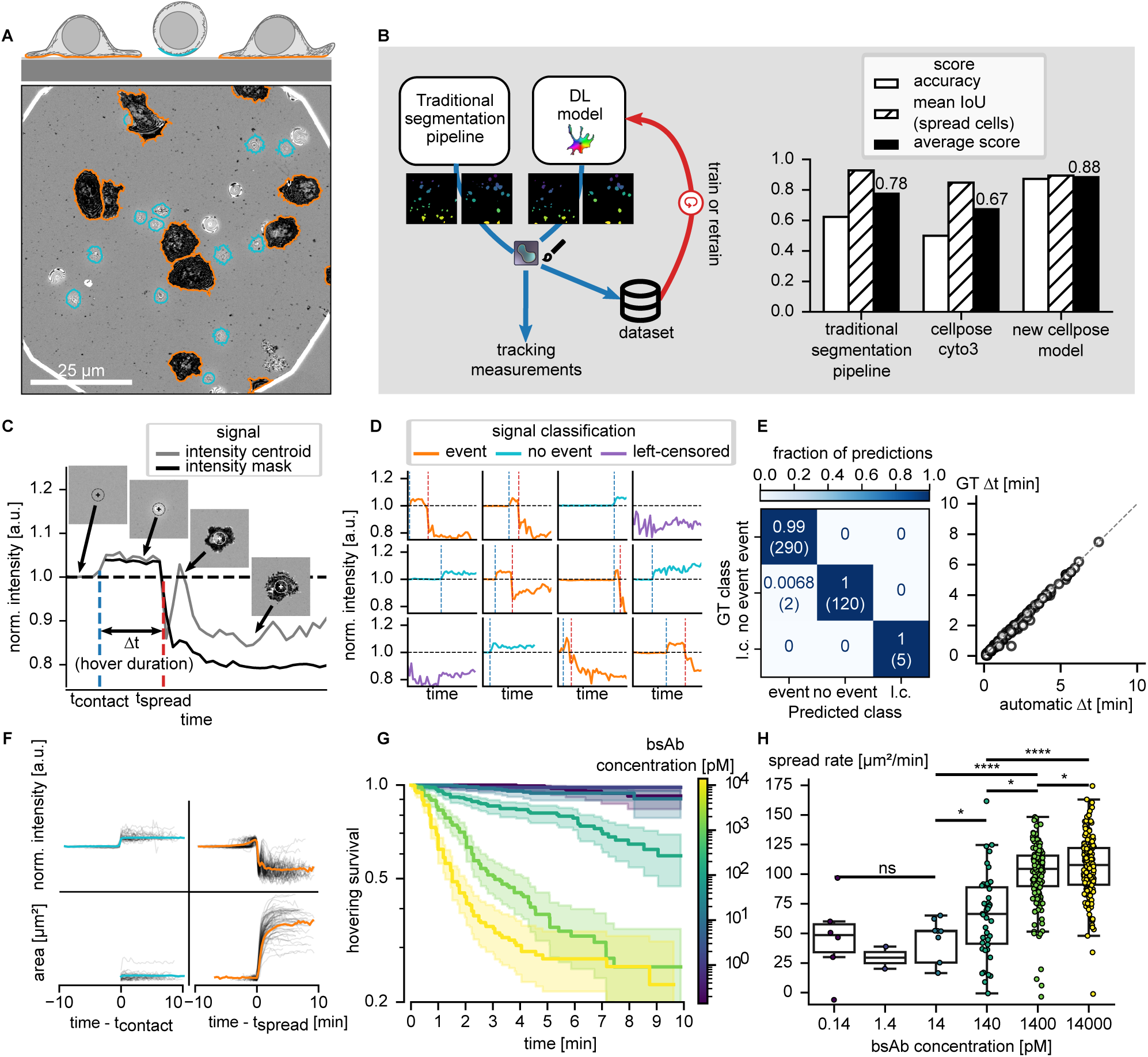
Functional response of immune cells in a spreading assay. A) Schematic (top) and snapshot (bottom) of a spreading assay imaged by time-lapse RICM. Primary NK cells sediment on an HER2-coated surface in the presence of bispecific anti-CD16 *×* HER2 antibodies. Cells touch the surface in a desynchronised manner and may spread after a stochastic hovering duration. The contours of the cells are represented in orange for cells classified as spread on the surface and blue for hovering cells (corresponding colours are also used in the schematic). B) A traditional segmentation pipeline is used to generate a first instance segmentation of the cells imaged in RICM. Cell masks can be manually curated before training a new Cellpose [37] model, either from scratch or by fine-tuning an existing generalist model. Barplot comparing segmentation scores obtained for three different methods that can be called within the software. With the improved results yielded by the new model, the user can proceed to tracking and measurements. C) Intensity time series for a cell performing a contact and spreading sequence (see text for details). D) Single-cell intensity time series colour-coded for three classes: 1) the cell spreads during the movie (’event’, orange), 2) the cell is not observed to spread during the movie (’no-event’, green) and 3) the cell is already spread at the beginning of the movie (’left-censored’, purple). E) Benchmarking of cell class automatic determination reported by a confusion matrix, and comparison of spreading times determined automatically. F) Individual cell time series (tonal, morphological, etc – grey traces) are synchronised with respect to a characteristic time to measure the average population response at the event time (coloured lines).G) Hovering survival curves for different bsAb concentrations: fraction of cells that have not yet spread as a function of time after surface contact. The shaded area represents the 95% confidence interval of the survival estimate. H) Spreading rate (dA/dt at *t*_spread_) for different bsAb concentrations, displayed as boxplots. The error bars correspond to the 95% confidence interval. Statistical significance was assessed using the Kolmogorov-Smirnov test, with P *>* 0.05: ns, P *<* 0.05: *, P *<* 0.01: **, P *<* 0.001: ***, P *<* 0.0001: **** .

Cell segmentation in RICM images is challenging because the cells can be locally brighter or darker than the background, causing direct thresholding techniques to fail [63]. To address this, previous approaches have adopted special techniques, such as background correction and applying a variance filter before thresholding [11, 64, 65], which, however, was often inadequate and required expert intervention. Our goal here was to train a DL model to segment lymphocytes spreading or hovering on the surface, without the need for expert corrections. The complete strategy – background subtraction (Supplementary Fig. S3B), generating an initial set of labels with the traditional segmentation pipeline (Supplementary Fig. S4), correcting errors in the viewer to build a training dataset (Supplementary Fig. S5), and training a DL model from this dataset – is illustrated in Figure 2A–B. Moving away from this application, Celldetective provides a whole ecosystem to perform segmentation and adapt deep learning methods (see Supplementary Fig. S6). Here, we trained a new Cellpose model on the RICM dataset (both the dataset and model are available on Zenodo). We benchmarked this model against the traditional segmentation pipeline and the *cyto3* generalist Cellpose model [38] using a test set, quantifying lymphocyte detection accuracy and segmentation quality separately (see Materials and Methods). Hovering cells, which appeared as bright blobs with interference patterns and had an ill-defined area in RICM, were excluded from the segmentation quality score. Our model achieved a higher detection accuracy (0.86), compared to the traditional pipeline (0.62) or the generalist model (0.5). The traditional pipeline tended to produce many false-positive detections, while the generalist model struggled to simultaneously detect spread and hovering cells. Segmentation quality, however, was higher for the pipeline (0.93) than for our model (0.89) or the generalist model (0.88). Since most spread cell labels in the test set required little manual correction (except for contiguous cells), we were not surprised to observe the highest segmentation quality with the pipeline method. The generalist model segmented spread cells well when it could detect them. Overall, the new Cellpose model achieved the best average performance across both tasks (0.88), making it the most effective approach for segmenting cells in this assay.

Next, tracking of cells — which move while hovering and are quasi-stationary while spreading — was performed using a versatile motion model (see Materials and Methods), without feature inputs to prevent sharp morpho-tonal transitions from disrupting tracking. We enabled track extrapolation from the beginning of the movies, allowing intensity measurements to start before the cells reached the surface (Figure 2C-D). With track post-processing, we could measure intensities prior to cell arrival, with a position-centred average intensity measurement, in a user-defined disk (position-based), which compared well with the average over the cell mask, when available (mask-based). Notably, the position-based intensity showed a spike during the spreading phase which is characteristic of the cell nucleus approaching the surface and that cannot be detected in the mask-based intensity.

We defined two characteristic event times (Fig. 2C): contact time (*t*_contact_)—the moment when the cell first appeared in RICM, with an average intensity greater than 1; and the onset of spreading (*t*_spread_) when the average intensity dropped below 1. The hovering time (*t*_spread_ − *t*_contact_), for spreading cells, is the time required to make a spreading decision. *t*_contact_ is automatically detected as the time of first segmentation. For *t*_spread_, spreading events were identified through condition-based classification (mask-based RICM intensity *<* 1) and an irreversible event hypothesis (see Materials and Methods). We benchmarked classification performance and hovering duration estimates against a manually annotated test set (Fig. 2D). The set was annotated with a dedicated viewer (see video ricm events.mp4). A confusion matrix indicates perfect agreement between automatic classification and ground truth, while a correlation plot comparing estimated hovering durations to the ground truth shows excellent agreement, with a mean absolute error of 0.35 minutes (Fig. 2E).

After automatically extracting the event times, we examined average population response based on single-cell time series, by synchronising each single-cell measurement to a specific event time to calculate the mean response across the population (Figure 2F). Next, we tested whether varying the concentration of bsAb on the surface impacts spreading. First, we estimated the “hovering survival” by plotting the fraction of cells hovering at a given time (Figure 2G). We observe a higher fraction of spreading cells and shorter spreading decision times with increasing bsAb concentrations. The acceleration of the spreading decision time was particularly apparent in the 10-1000 pM bsAb concentration range, saturating beyond 100 pM. Next, using the table exploration feature in Celldetective, we differentiated the area with respect to time for each cell track, extracting the spreading rate at the onset of spreading (*t*_spread_), which increases in the 14-1400 pM range (Figure 2H). Overall, determining event times for individual cells, measured at high throughput, highlights Celldetective capabilities in evaluating cell-surface interactions.

### Cytotoxic response in a co-culture with overlapping channels

In the second application, we further tested the capacity of the same bsAbs to induce cell-cell contacts and trigger antibody-dependent cell cytotoxicity (ADCC). For this purpose, we used a high-density co-culture of target tumour cells (MCF-7) and effector cells (primary NK). The concentration of anti-CD16 × HER2 bsAb, as well as tumour cell HER2 expression, are varied. The lysis of target cells is monitored by Propidium iodide. The assay also includes a fluorescent label of the live target cell nucleus as well as the effector cell cytoplasm, in order to locate both cell types over time. A multi-colour composite representation of a single position is shown at endpoints in Figure 3A, illustrating how the number of dead (red) nuclei increases over time.

**Fig. 3.**
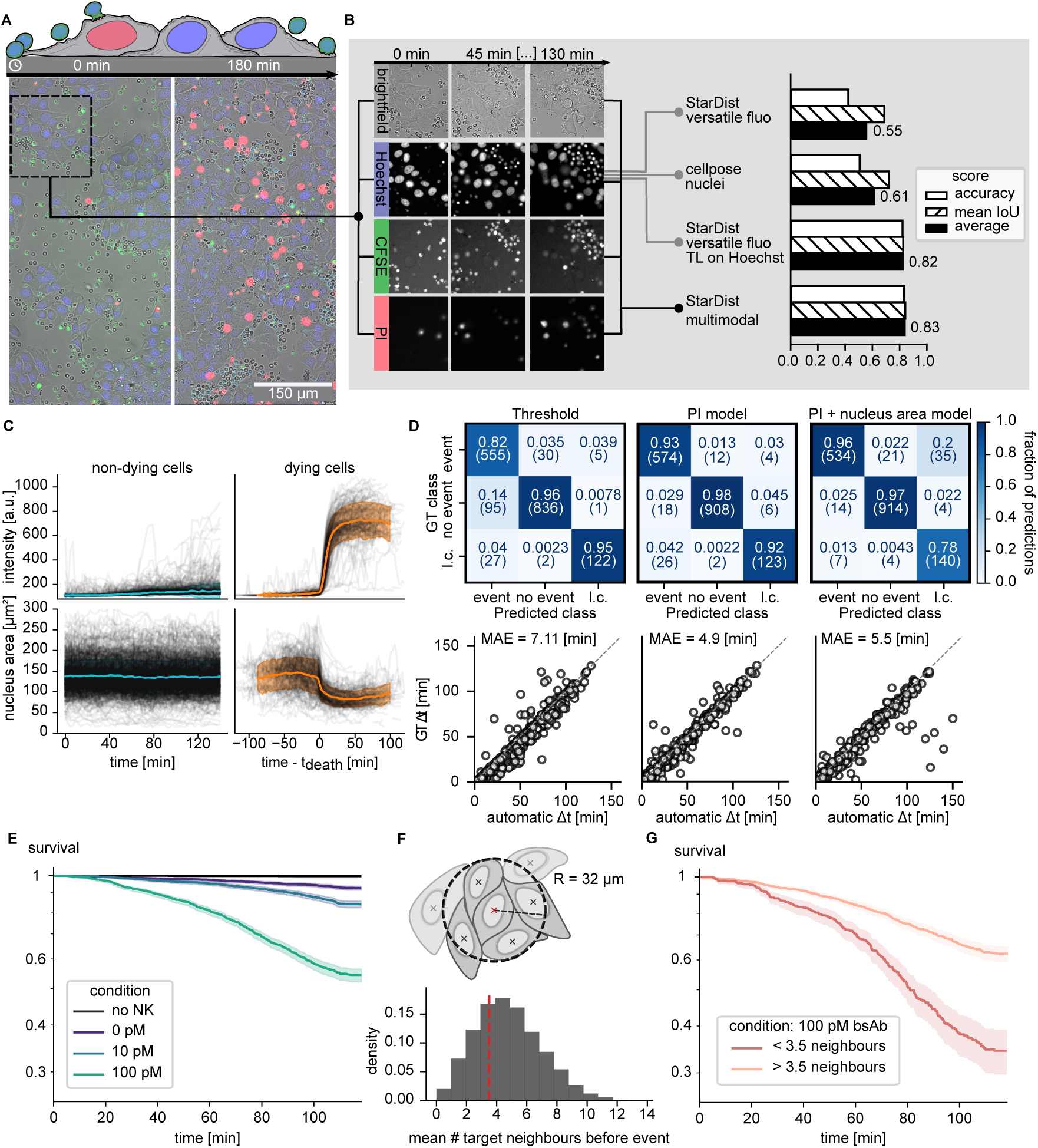
High-throughput cytotoxic response of cancer cells in co-culture with immune cells. A. Schematic side-view of target/NK cells co-culture assay for bispecific ADCC (top) and representative multimodal composite images, obtained at two different time points, with target nuclei labelled in blue, dying cells in red and NK cells in green (bottom). Corresponding colours are also used in the schematics. B. Decomposition of partly overlapping fluorescence channels and benchmark of segmentation DL models. Brightfield and fluorescence images at three different time points. Hoechst channel (target nucleus) is taken as input to the existing StarDist *versatile fluo* or Cellpose *nuclei* models (available directly in Celldetective). Two new models trained in Celldetective were benchmarked: a transfer of the StarDist *versatile fluo* model on our MCF-7 nuclei dataset, with Hoechst as input, and a StarDist multimodal model trained from scratch using 4 channels as its input. C. Time series of nuclear fluorescence intensities and nucleus apparent area for a set of non-dying target cells (left) and of dying cells (right); the reference time *t*_0_ is the death time for the dying cells. Individual traces in black, average in colour. The coloured shaded area the standard deviation. D. Benchmark of three methods for event classification and regression (Threshold method on PI, DL classifier on PI, DL classifier on PI and nuclear area). Top row: confusion matrices with 3 classes (fraction of predictions displayed). Bottom row: correlation plots for the precision on death time determination. E. Survival curves of target cells without NK cells or in the presence of NK cells for different bsAb concentrations. The shaded area represents the 95% confidence interval of the survival estimate. F. Schematics for neighbour counting and histogram of target neighbour counts (time-averaged per cell). The dashed line at 3.5 neighbours represents the median of the distribution, used to split the population into low- and high-density subgroups. G. Survival curves for two subpopulations of targets at 100 pM bsAb, split as a function of the local target cell density (below or above 3.5 neighbours). The shaded area represents the 95% confidence interval of the survival estimate. The non-integer threshold of 3.5 arises because the neighbour count is time-averaged over the observation period for each cell.

First, we studied the lysis behaviour of target cells over time, which entailed achieving a single-cell description for the target cells. Directly using generalist nuclear segmentation models (StarDist *versatile fluorescence* and Cellpose *nuclei* models) on the Hoechst channel, we could only achieve a poor accuracy, limited by occlusion of the underlying target cells by effector cells. While it was possible to segment all cells and classify them either simultaneously [66] or in two steps [67], label contamination from overlapping populations remained an issue. We produced expert annotations of the MCF-7 nuclei using all available multi-channel and time information. Since the MCF-7 nuclei are larger than primary NK cell nuclei, we expected to be able to train an efficient model detecting the MCF-7 nuclei from the Hoechst channel only, failing only on blurry images, where the size separation was no longer obvious (Fig. 3B). We used Celldetective to retrain the StarDist *versatile fluorescence* model on our dataset using the Hoechst channel as its input and considerably improved the performance, from 0.55 to 0.82. Last, we trained a completely multi-channel model on all four available channels to detect the MCF-7 nuclei and achieve a slightly higher performance (0.83). All of the models benchmarked are accessible directly in Celldetective (see Supplementary Tab. S1 and S2). Target cell tracking is detailed in Materials and Methods.

An expert annotated the death events, defined as a sharp increase in PI signal at *t*_death_. Single-cell time series were then grouped by event class and synchronised to *t*_death_ (Figure 3C). The average PI intensity for cells undergoing death shows a distinctive step-like transition with low lateral variance, validating the annotation accuracy (Fig 3C, top-right). The PI signal in live cells, by contrast, rises slightly over time but remains well below the levels observed in dead cells (Fig 3C, top-left). Notably, the apparent nuclear area of dying cells contracts sharply at *t*_death_ (Fig 3C, bottom-right), while the nuclear area of non-dying cells remains constant (Fig 3C, bottom-left).

Event detection was evaluated for three methods on this dataset: (1) a condition-based classification using the irreversible event hypothesis on the PI intensity time series, (2) a DL model trained to detect death events from PI alone, and (3) a DL model trained to detect events using both the PI intensity and nuclear area time series. Detailed information on the training procedures for these models is provided in the Materials and Methods section and the DL architecture on Fig. S7. Figure 3D presents the confusion matrices for each method, showing agreement between predicted and ground-truth event classes, along with a correlation plot evaluating the accuracy of death time extraction. Method 1 incorrectly predicted about 15 % of the events as death events when none had occurred. This method also had the highest mean absolute error (MAE) for death time estimates (7.1 minutes). Method 2 performed better, with only 3 % of its predicted death events corresponding to an absence of events. The multivariate model (Method 3), incorporating both PI and nuclear area data, achieved the highest precision in death event detection (96 % of predicted death events were true events). However, this model showed a reduced accuracy in identifying cells as already dead, with 20 % of these cells (2 % of the total number of cells) observed to die later. This remains a minor drawback here, since the fraction of initially dead cells is low. Celldetective’s event annotation viewer allows the user to efficiently review and manually reclassify these cells, recovering them for the survival study. As before, quality check and manual corrections were performed with the event annotation viewer, using a multichannel composite of the images (Video adcc rgb.mp4).

We next investigated whether the probability of tumour cell lysis depended on bsAb concentration. Using *t* = 0 min as the synchronisation time, we excluded cells already dead at the start (always less than 5%) and disregarded cells born from mitosis (during tracking post-processing). We represented the MCF-7 survival curve using *t*_death_ as the event time (Figure 3E), which shows an increase in the fraction of dead MCF-7 cells (endpoint) occurring at a faster rate with rising bsAb concentrations.

Our single-cell approach also permits us to link cell survival with local context. To do this, we calculated a target-target neighbourhood within a 32 *µ*m radius (chosen to encompass approximately two cell diameters – MCF-7 nuclear diameter ≈ 15 *µ*m), thus capturing the nearest layer of neighbours while excluding more distant cells, (isotropic method, see Materials and Methods and Fig. S8) to count the local density experienced by each target cell until death (Figure 3F). We then applied a filter in Celldetective to compare survival outcomes for cells in low-density versus high-density environments (Fig 3G), observing a significant survival advantage for targets in high-density regions. This effect was present at all tested bsAb concentrations, though less pronounced at lower concentrations where the overall killing rate was reduced.

### From a single time point to a dynamic readout of effector-target interactions

After describing the ADCC assay from the targets’ perspective, we shifted our focus to the effectors. The aim here was to characterise how effector cells interact with their target neighbours: first, we focus on the analysis of the distribution of fluorescently labelled bsAbs at the cell surfaces at the population level using instantaneous measurements; second, we study the relative motion between targets and effectors in relation to a degranulation markers using cell tracking and pair analysis. Effector cells were segmented using a custom multichannel Cellpose model that combines brightfield, Hoechst, and CFSE, a cytoplasmic fluorescence label (see Supplementary Tab. S3). We chose Cellpose for effectors because NK cells have irregular, non-convex shapes, for which the Cellpose algorithm is better suited, while StarDist, which assumes star-convex objects, was more appropriate for the roughly elliptical MCF-7 nuclei segmented in the previous section. Unlike targets, effectors exhibit high motility and can form dense clusters in the assay, limiting reliable tracking.

When effector tracking was unfeasible (e.g., at an effector-to-target ratio of 5:1 and frame interval greater than 2 minutes), we could still perform a multicondition analysis of the cells at each time point. We aimed to measure the signal of antibodies tagged with a single ATTO 647 dye. Specifically, we compared the CE4-21 bsAb with the CE4-X control antibody, which only binds to the HER2 antigen on targets, across two target cell lines with distinct antigen expression levels (WT and HER2+) [68]. Figure 4 presents the mean bsAb fluorescence intensity under effector cell masks at 1 hour, broken down by target cell type, antibody, and contact state (in or off contact with targets, see Materials and Methods and video single-timepoint-contact.mp4). For WT targets (lowest HER2 expression), the cells exhibited a signal in the CE4-21 bsAb condition, regardless of the contact state (Figure 4A). In contrast, no signal was measured with the CE4-X antibody condition, supporting that the bsAb-condition signal came from the effector cells (visual assessment in Figure 4B). With the HER2+ targets and CE4-X antibody, the signal measured off-contact was negligible, confirming that the antibody did not bind to the effectors (Fig 4A-B). The signal measured in contact did not originate from the effector cells but from the target cells below. With the CE4-21 bsAb, on the other hand, the signal measured in contact was a sum of contributions from both the target and effector cells.

**Fig. 4.**
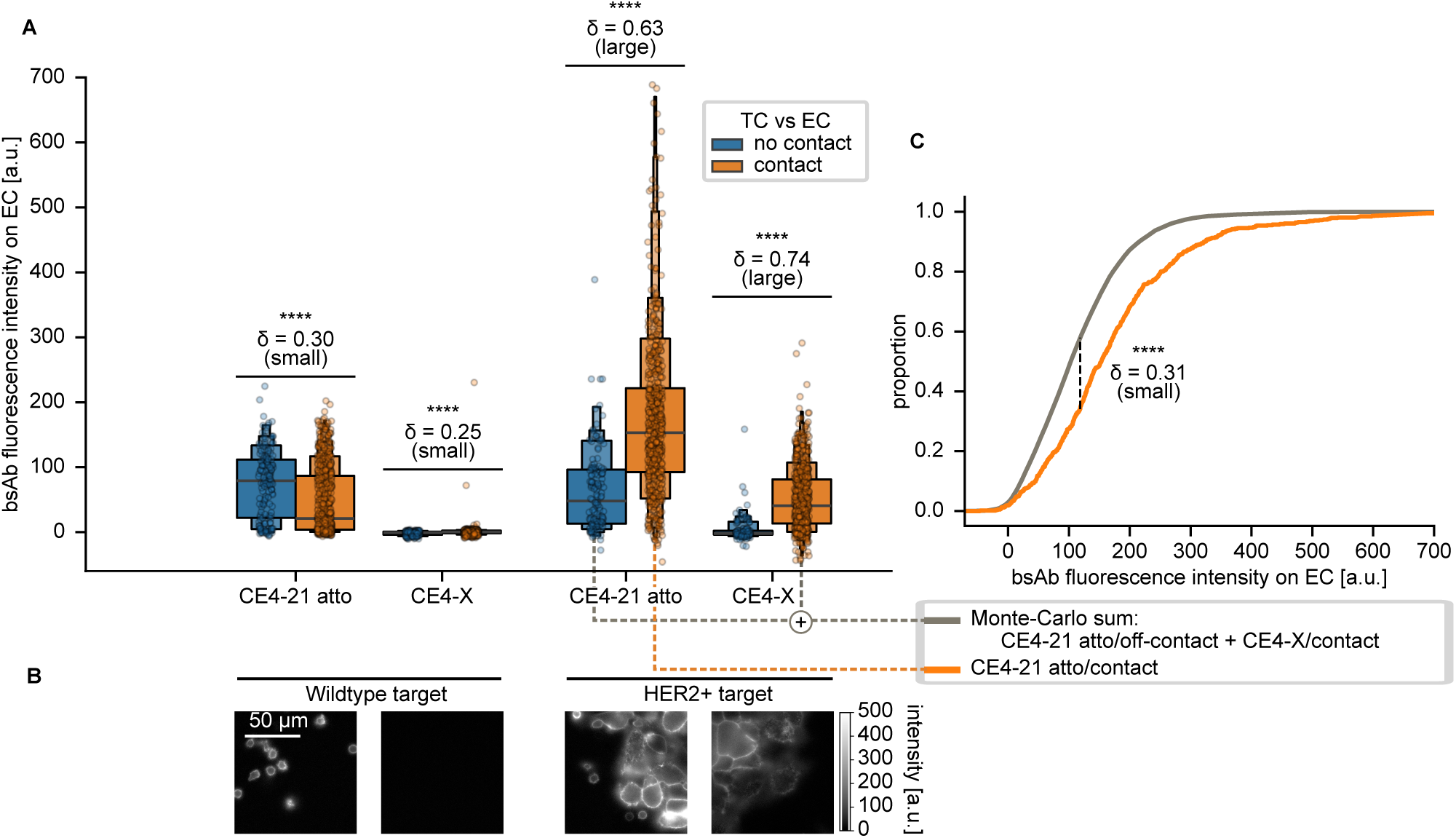
Single time point analysis of effector-target interactions. In the absence of effector tracking, fluorescent bsAbs were used to study the distribution of Ab on effectors and targets. A. bsAb intensity measured on effector cells on targets with high or low antigen expression. bsAb CE4-21 binds to both HER2 and CD16, while CE4-X control antibody binds only to the HER2 antigen on target cells. B. Representative snapshot of fluorescence channel for each condition of A. C. Empirical cumulative distribution functions (ECDFs) for the simulated (grey, sum of CE4-21/no contact + CE4-X/contact) and observed (orange, CE4-21/contact) distributions. The dotted black line represents the Kolmogorov-Smirnov statistic, *i.e.* the maximum separation between the two ECDFs.

Based on the measured signals, we sought to determine whether the added effector signal from the CE4-21/off-contact condition (without target contribution) and the effector signal from the CE4-X/in-contact condition (without effector contribution) could approximate the CE4-21/in-contact signal. Using the distributions generated by the software, we simulated the distribution of this summed signal with a Monte-Carlo approach, and compared it with the CE4-21/in-contact condition. Figure 4C reveals that the simulated signal distribution was significantly lower than that of the CE4-21/in-contact condition, with a small effect size (Cliff’s Delta *>* 0.33). This excess signal suggests that, upon direct effector-target contact, the bispecific antibody is enriched at the cell-cell interface through simultaneous engagement of CD16 on the NK cell and HER2 on the target, providing direct image-based evidence of bsAb bridging at the immunological synapse. By calculating this effect size at each time point, we further confirmed the robustness of this finding, as the effect consistently maintained the same direction, with a medium effect size observed more often than not (Suppl. Fig. S9).

If effector tracking is achievable (e.g., at an effector-to-target ratio of 1:1 and frame interval lower than 1 minute), we can explore the measurement time series. To monitor the lytic activity of effector cells dynamically, we added a degranulation marker (fluorescent anti-LAMP1), exploiting Celldetective’s multivariate time series capabilities. In this experiment, we compared two bsAbs, CE4-21 and CE4-28, formed of the same CE4 nanobody against HER2 and two different nanobodies against CD16 on the effector side. An example of such labelling is shown in Fig. S10.

First, we investigated whether LAMP1 expression was influenced by biological conditions, mirroring our approach with fluorescent antibodies. For each target cell type and antibody condition, we reported the fraction of LAMP1-positive effectors, decomposed by contact state, across all time points (Fig. 5A). The results indicated that, except for the no-antibody controls, the fraction of LAMP1-positive effector cells was consistently higher in-contact with target cells.

**Fig. 5.**
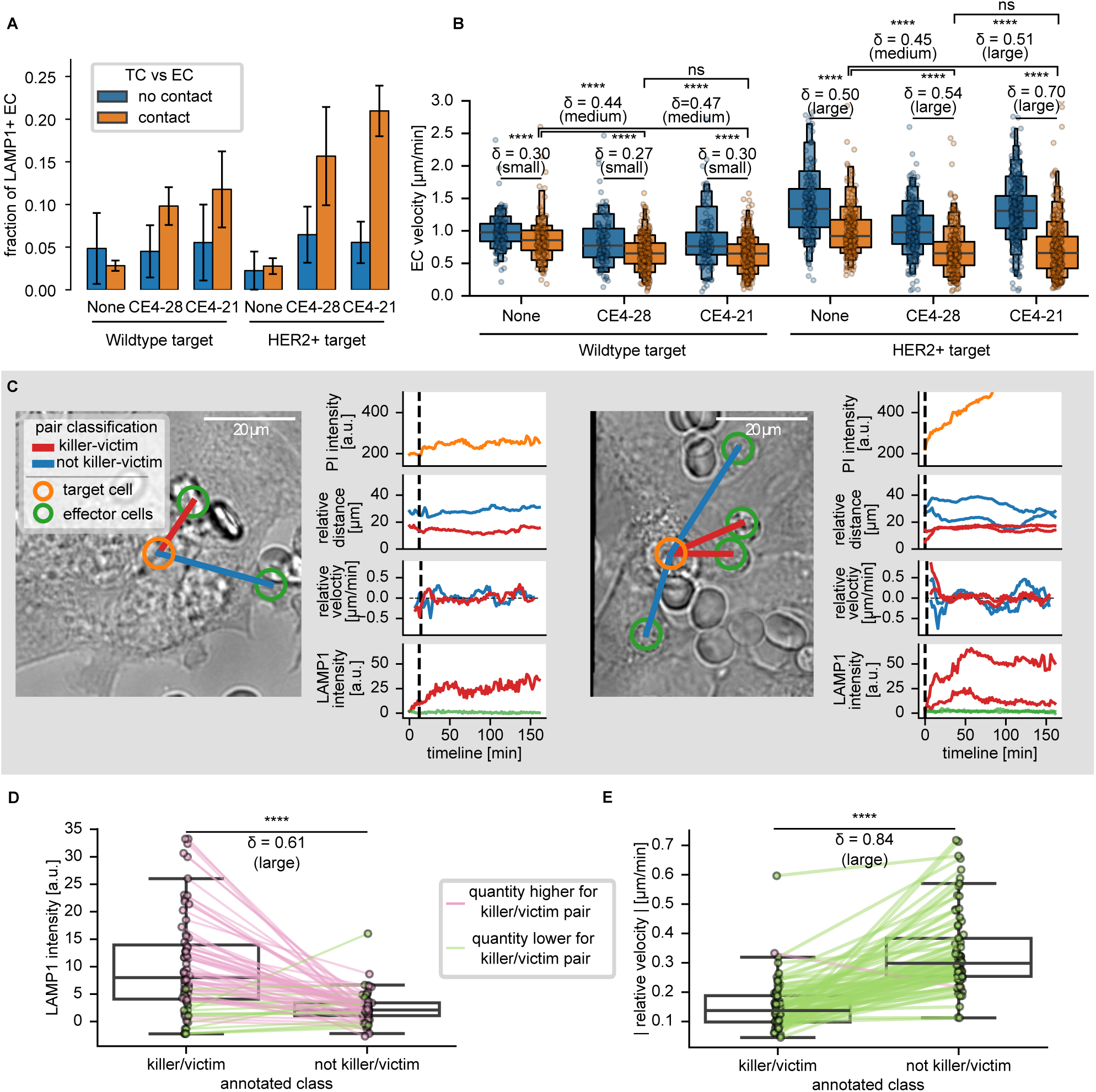
Effector-target dynamic interactions. A. Fraction of LAMP1-positive effector cells for different bsAb conditions, decomposed by contact with the targets (over all time points). Bars indicate the mean fraction across three replicate positions; error bars denote the standard deviation. B. Average effector velocity for different bsAb conditions, decomposed by target contact (up to two estimates per effector). C. Examples of killer-victim identification with one (left) or two (right) killers. One target cell (centre) is connected to effector neighbours via coloured segments. Dynamic effector-target interaction monitoring includes PI signal of target cell, relative distance and relative velocity between target and effector, as well as LAMP1 signal in effector neighbours. D, E. Characteristic parameters for killer/victim and non-killer/victim pairs. Effector cells are manually annotated as potential killer or not. Each pair of points represents one victim target, with the average over all neighbours of the parameter decomposed by killer class. D. LAMP1 intensity. E. Relative effector-target velocity.

From this time-integrated approach, we shifted our focus to the dynamics of the effector cells, specifically investigating whether their motility was influenced by target cell proximity. Given that velocity estimators can be noisy, we computed a time-averaged velocity estimate for each cell track, decomposed by contact state. The resulting velocities for all conditions are presented in Figure 5B. In each condition, the contact velocities were consistently lower than off-contact, regardless of antibody presence or type. This difference was always significant, with a small effect size for WT targets (Cliff’s Delta *>* 0.147) and a large effect size for HER2+ targets (Cliff’s Delta *>* 0.474). The lowest contact velocities were observed in the presence of antibodies, showing a significant drop from the target-specific control, with medium effect sizes for the WT bsAbs and HER2+ CE4-28, and a large effect size for HER2+ CE4-21. However, we did not detect any difference between the two antibodies, separately for each cell type. We note that off-contact velocities appear slightly higher on HER2+ targets across all conditions; this is likely due to differences in target cell monolayer coverage and density between wells, which can influence effector motility independently of the antibody condition.

After computing effector-target neighbourhoods, we measured features of cell pairs, including relative distance and angle, as well as their time derivatives. Figure 5C illustrates time series data for pairs, showing the PI intensity of a target cell that died at 5 minutes alongside the relative distance, velocity, and LAMP1 signal of neighbouring effectors. Notably, only the selected neighbour (highlighted in red) exhibits a strong LAMP1 signal, which rises around *t*_death_, consistent with degranulation preceding or coinciding with target membrane permeabilisation, and remains elevated until the end of the observation.

Using the pair annotation feature, we aimed to identify “killer-victim” pairs. An expert annotated each neighbourhood containing a dead target cell, identifying the most probable effector cell killer to establish these pairs. This annotation was done via a cell pair viewer, similar to the viewer designed for single cells, which leveraged colour cues for pre-determined target cell death events and immobilised effector cells (Video cell interactions.mp4). The expert viewed only the brightfield movie, blinded to the LAMP1 signal, and identified effectors with prolonged, restricted movement in contact with the target cell. Effectors that arrived post-mortem were excluded, and in ambiguous cases, multiple pairs could be labelled as “killer-victim” within a neighbourhood. Due to the complexity of this task, precise event times were not annotated. Figures 5D-E display average relative velocity and LAMP1 expression for each neighbourhood, with values grouped by killer-victim status and shown in paired boxplots. Results indicate significantly higher LAMP1 expression and lower relative velocity in killer-victim pairs, with both metrics showing a large effect size (Cliff’s Delta *>* 0.474).

## Discussion

Here, we developed a new software tool to study multichannel time-lapse microscopy data at scales ranging from single cells to cell populations. It integrates original methods for event detection and cell-pair analysis with established methods for cell segmentation and tracking [36, 37, 42]. Although the pipeline steps are primarily sequential (see Fig. 1, S2) and could be performed independently with different tools and scripts, we demonstrate how integrating them within a unified, interactive software environment enhances data exploration and facilitates the analysis of dynamic cell interactions in large datasets.

Celldetective demonstrated its capacity to manage datasets of hundreds of gigabytes, generating high-quality plots that include hundreds of single-cell traces and data points. As a standalone application, it empowers biology experts to analyse single-cell data, annotate labels and events, and train models to automate tasks without requiring coding skills—similar to recent advances for fixed images [55]. Celldetective integrates the open-source annotation tool napari for label correction and introduces unique tools for event annotation and pair classification. The software combines traditional threshold-based image and signal analysis methods with deep learning-based approaches, which can be used complementarily for enhanced results.

We showed how recording multivariate time series may reveal multiple phenomena occurring during the same event. For example, cell death, monitored through an increase in PI fluorescence, was observed alongside apparent nuclear shrinkage (see Fig. 3C). We interpret this as cell de-adhesion upon death, releasing the pushing pressure on the nucleus and allowing its axial shape relaxation [69]. In immunotherapy assays involving human primary cells, we prioritised a fluorescence-based labelling strategy that avoids genetic modification. Due to the exchange of fluorescent dyes between cell populations and cell lysis during the assay, along with significant size disparities between effector and target cells, generalist deep learning segmentation models perform poorly. To address this, we trained custom models within the software, using StarDist for target nuclei segmentation (convex object) and Cellpose for effector cells (any shape). Performance improved slightly with the simultaneous use of multiple input channels (see Fig. 3B). On closer inspection, the improvement was more noticeable on difficult data (blurry, noisy). This approach also benefited event detection: training a model on both PI and nuclear size time series enhanced the event classification performance on the two most important classes for survival studies (see Fig. 3D). Our spatially resolved immunotherapy assay documents the effect of heterogeneity in target cell density. Indeed, we observed how the high density of target cells reduces their ADCC killing rate (see Fig. 3G). This survival advantage in dense regions could reflect a direct protective effect of close target neighbours (for instance through physical shielding or competition for effector engagement) or alternatively a sampling effect, where effectors in high-density areas are diluted across more targets, reducing the per-target encounter rate. Disentangling these mechanisms would require further experiments with controlled cell densities, which is beyond the scope of this software-focused study.

Immune cells leverage transient and weak contacts to make rapid decisions [10]. We demonstrate the power of high-throughput imaging and analysis to explore such mechanisms. In a cell-surface assay, we measured by interference microscopy the decision time for NK cells to spread on an antigen surface in the presence of bispecific antibody. Celldetective provides a higher statistical power than in earlier works [61]. Thus, we observed differences in NK response to bispecific antibody concentration in the minute range, which could not be resolved in earlier studies on similar systems such as TCR-pMHC [70], where the smaller cell numbers and lower throughput limited the statistical power. Celldetective’s high-throughput capabilities could in principle help resolve such subtle kinetic differences in T cell assays, provided the experimental conditions (e.g., imaging modality and cell density) permit reliable segmentation and tracking. Here, the detection of sharp cell morphology changes during adhesion and spreading could be further enhanced by combining RICM with brightfield imaging, since each imaging modality exhibits a contrast specific to each cell state [60].

Proximity between effector and target provides spatial information inaccessible to traditional flow cytometry, revealing clear differences in LAMP1 signal and relative velocity in neighbourhoods of targets. We introduce a classification scheme, which allows quantification of a subpopulation based on arbitrary thresholds on any measurement. Additionally, instantaneous classification can be complemented by a time series analysis, when cell tracks are available.To extend this analysis to effector-target dynamic interactions, we addressed the question of killer:victim pair identification. The strategy consisted of collecting relative measurements like velocity and manually annotating the potential killer for each target from the images. Our data are consistent with one or more killers. The most prominent parameters for killer identification are relative velocity and LAMP1 signal. Since cell pairs were implemented like single cells, Celldetective allows users to train an event detection model from pair, target and effector time series to automate killer:victim pair identification. As a future direction, further work is required to automatically analyse the dynamic signalling and provide more robust prediction. Celldetective offers in principle the complete environment for such an advanced analysis, as demonstrated by the automatic detection of death times.

For its applicability, Celldetective does not require coding skills. It was tested by graduate students from our labs with diverse backgrounds in biophysics, cell biology and immunology. Its release is accompanied by worked examples, tutorials, and comprehensive documentation structured around the Díataxis framework (tutorials, how-to guides, explanations, and reference; see https://diataxis.fr/). Software maintenance and user assistance are offered via the GitHub platform and a professional development team (https://github.com/centuri-engineering). One highlight of Celldetective is to offer alternative methods, in particular for cell segmentation and tracking, so that the user can maximise the outcome. In order to guide the user for software applicability and workflow, we propose a decision tree, visible as question marks in the interface, to assist in choosing between the different options.

The experimental systems analysed here are largely two-dimensional, but at the cellular scale, exhibit a three-dimensional aspect. Celldetective is specifically optimised for high-throughput, high-temporal-resolution imaging of quasi-2D systems. By prioritising temporal sampling over Z-axis depth, the software enables the capture of rapid biological dynamics that are often the focal point of interaction studies. Future software expansions could incorporate 2.5D or 3D sequence analysis, facilitated by the use of 3D-compatible backends (StarDist, Cellpose, bTrack). Currently, to our knowledge, only proprietary solutions are offered and exploited in immunotherapy assays [27]. Notably, cell-interaction studies benefit from higher time resolution, which may limit compatibility with extensive 3D analysis.

The methodologies we demonstrate here should adapt well to commercial high-throughput automated microscopes for long-term cell co-culture studies [40], for which no efficient analysis solutions are available, to our knowledge. A distinctive aspect of our approach is the use of compact, efficient models that are user-trainable on a single GPU setup. This contrasts with approaches relying on large-scale databases [40] and extensive DL models [71]. Moreover, the software’s integrated visualisation and annotation features encourage meticulous review of segmentation and tracking results, enhancing confidence in the analysis. Convolutional models used by the software, such as U-Net and ResNet, are prone to forgetting previously learned tasks upon transfer [72]. To address this, Celldetective allows the addition of the original dataset during fine-tuning from a previously trained model. Future extensions could include foundation models like Segment Anything [73] for broader adaptability.

## Conclusion

In conclusion, Celldetective is a user-friendly software tool designed for non-specialists, providing advanced cell-scale analysis for large cell populations. Combining high time-resolution with robust statistical capabilities, we show its potential to uncover new fundamental parameters characterising dynamic interactions between immune and tumour cells. We demonstrated the efficiency of Celldetective through two applications in the context of immunotherapy by isolating time-dependent cell-scale parameters and pair interactions, in a high cell-density environment. We believe that such an approach holds great potential for enhancing the design and discovery of drugs in a more powerful and rational manner, while its general design suggests potential for broader application to other biological contexts where dynamic cell-cell interactions are of interest.

## Materials and Methods

### Software and hardware

The software, implemented as a Python package with a PyQt5 graphical user interface (GUI), incorporates Material Design for styling and integrates Matplotlib canvases for displaying images, animations, and plots. Each processing module—segmentation, tracking, measurement, event detection, and pair analysis—operates on a single movie at a time within an independent subprocess. Upon completion, CPU and GPU memory are fully released, enabling seamless iteration on the next movie. Where applicable, multi-threading has been implemented and can be customised within the software. Each module generates outputs (images or tables) saved directly to the experiment folder, ensuring data preservation and modularity. This design allows users to import data flexibly from external tools, including ImageJ and Cellpose UI. Detailed outputs include mask images from segmentation, trajectory tables from tracking, and additional columns for measurements and event detection. The neighbourhood module further extends the trajectory tables, producing a pickle file with a nested-dictionary structure that logs neighbour information at each time point.

Development and testing were conducted on various hardware configurations: an Intel Core i9 CPU, NVIDIA RTX 3070 GPU, and 16 GB RAM desktop running Ubuntu 20.04 for primary development; extensive testing was completed on both Windows 11 (Intel Core i9 and Core i7 laptops) and an older Ubuntu 20.04 desktop with an Intel Core i5 CPU. Celldetective is routinely validated across Python versions 3.9 to 3.11 on both Windows and Linux, with a comprehensive automated test suite of 43 test files (26 GUI-level and 17 unit tests) covering segmentation, tracking, measurements, event detection, viewers, and table operations, integrated with continuous integration and deployment (CI/CD) on every commit.

### Antibody design and production

The bispecific antibody (bsAb) C7b-21 (or CE4-21, CE4-28) is a fusion of two single domain antibodies (sdAb, also called nanobodies), sdAb C7b (or CE4) targeting human epidermal growth factor receptor-2 (HER2/neu or ErbB2) [74] and sdAb C21 (or C28) targeting human CD16 (Fc*γ*RIII), using the human CH1/Ck heterodimerisation motif, corresponding to the bispecific Fab (bsFab) format [75]. bsFab CE4-28 was produced by co-transfection of two plasmids using the mammalian transient system Expi293 (Thermofisher) and purified as previously described [76]. For experiments in Fig. 4, bsAb CE4-21 and control antibody CE4-X were specifically labelled with Atto647 dye using transglutaminase reaction (Zedira, Darmstadt, Germany).

### Cell lines culture, effector extraction and phenotyping

Modified MCF7-HER2+ cells were stably transfected from the MCF7 cell line to overexpress HER2 receptors. Determination of HER2 levels on the cells was performed by flow cytometry with Herceptin antibody and a secondary fluorescent anti-human antibody. Fluorescence intensity was determined and correlated with HER2 levels. Cells were maintained in RPMI 1640 medium (Gibco, Life Technologies), supplemented with 10% Fetal Bovine Serum, FBS (Gibco, Life Technologies), 50 mg/mL of hygromycin for antibiotic selection. Cells were split three times per week and kept in the incubator at 37*^◦^* C under 5% CO_2_ atmosphere.

Primary human NK cells were isolated as described in [60]. Briefly, blood samples were obtained from the Établissement Français du Sang (Marseille, France), using the MACSxpress whole-blood human NK cell isolation kit (Miltenyi Biotec, Bergisch Gladbach, Germany); a negative selection was performed. The characterisation of the sorted cells was determined by flow cytometry with anti-CD16, anti-CD3 and anti-CD56 antibodies. Cells were maintained in RPMI 1640 medium and 10% FBS at 37*^◦^* C, 5% CO_2_ and used within the following 24 hours.

### Cell-surface assay

Primary human NK cells, freshly isolated and prepared at a concentration of 200,000 cells/mL, were introduced into an uncoated eight-well chambered polymer coverslip-bottom (Ibidi, 80821). For surface preparation, each well was initially rinsed with PBS, then incubated with 100 *µ*g/mL Biotin-labelled bovine serum albumin (BSA; Sigma A8549-10MG) in PBS for 30 minutes at room temperature (RT) with agitation. Non-adsorbed BSA-biotin was removed by four PBS rinses. Next, a 10 *µ*g/mL Neutravidin solution (Thermo Scientific, 31000) in PBS was added for 30 minutes at RT with agitation, followed by another four PBS rinses. The wells were then incubated with a 10 nM solution of Her2/ERBB2 protein (Sinobiological, 10004-H27H-B) in 0.2% BSA for 30 minutes at RT with agitation and subsequently washed four times with PBS. After this surface preparation, the Ibidi chamber was placed under a microscope at 37*^◦^* C. Prior to NK cell introduction, the cells were pre-incubated with bsFab (C7b-21) at the desired concentration for 30 minutes at 37*^◦^* C. They were then injected into the prepared sample.

Cell-surface contact and spreading dynamics were monitored using reflection interference contrast microscopy (RICM), which is sensitive to variations in cell-surface distance. The RICM configuration, described in detail by Limozin et al. [63], employed an antiflex Zeiss objective (NA = 1.25) and a green LED light source (*λ* = 546 nm) with a 14-bit CCD detector (Andor iXonEM, Oxford Instruments). This setup allowed for live cell observation at 37*^◦^* C. Images were taken after cell deposition in the chamber, with sixteen fields of view selected per condition and monitored through cyclic imaging over a 10-minute interval. The motorised stage (Physik Instrumente, Germany) facilitated sequential imaging, with temporal intervals between images in each field ranging from 15 to 20 seconds, enabling analysis of decision time between cell landing and spreading as well as NK cell spreading kinetics.

### Cell-cell assay

MCF7-Mod cells, expressing HER2 at a surface density of approximately 500 molecules/*µ*m^2^ [68], were used as targets. 80,000 cells per well were seeded into an 8-well chamber *µ*-Slide (Ibidi—Munich, Germany, polymer bottom, TC treated) and cultured overnight in RPMI supplemented medium under standard conditions (37*^◦^* C, 5% CO_2_) to reach exponential growth. The next day, the medium was removed, and cells were incubated with Hoechst 33342 (5 *µ*g/mL) in colourless RPMI at 37*^◦^* C for 10 minutes. After three rinses in warm RPMI with 10% FBS, cells were returned to the incubator.

Primary NK cells were labelled with CFSE dye following manufacturer instructions (CellTrace CFSE, ThermoFisher). 2.5 million cells were centrifuged at 1500 rpm for 5 minutes, supernatant was discarded, and cells were incubated with 2.5 mL CFSE in PBS for 20 minutes at 37*^◦^* C. Following a brief incubation in RPMI with 10% FBS (12.5 mL added to the cells for 5 minutes), cells were centrifuged, resuspended in complete medium, and incubated.

Dilutions of the bispecific antibody (bsAb) CE4-28 were prepared for final concentrations of 1, 10, and 100 pM. The bsAb solution (2 *µ*g/mL) was added to wells with Hoechst-labelled target cells, incubated at 37*^◦^* C for 20 minutes, followed by Propidium Iodide (PI; Sigma). NK cells were then added at an effector-to-target (E:T) ratio of 2.5, and the slide was placed on a Zeiss AxioObserver microscope with temperature control at 37*^◦^* C. Imaging was performed with a 20x/0.4 objective (pixel size 0.31 *µ*m) in transmitted light (brightfield) and epifluorescence channels (CFSE, Hoechst, PI), capturing 5-9 fields per well over 2-3 hours with 3-minute intervals between frames.

For NK cell tracking, observations were conducted using a 40x/1.3 Oil DIC (UV) objective (pixel size 0.157 *µ*m) with a 1.73-minute interval per frame, and the E:T ratio was adjusted to 1:1. To monitor degranulation, anti-LAMP1 APC-labelled antibodies (BioLegend, Clone H4A3, Cat. 328619) were added at 5 *µ*L per 1 million NK cells.

### Software methods

#### Tracking tuning

Celldetective provides graphical interfaces for two tracking backends: the Bayesian tracker bTrack [42] and the trackpy linking algorithm [59]. In the applications presented here, we used bTrack. We developed a configuration interface that enables users to view and adjust the bTrack configuration file (defining the motion model and tracklet reconnection parameters) or to import configurations from the bTrack-napari plugin. Typical bTrack configuration files are provided for the applications we present. Additionally, users can incorporate mask-based measurements to enhance the Bayesian update, with feature normalisation handled automatically by the software. Finally, users can set options for track post-processing (interpolation, extrapolation, filtering) to refine and correct the tracking data. A dedicated tracking correction module opens the segmentation masks in napari relabeled by track ID, allowing users to propagate a track ID onto any mask at any frame through a built-in interaction that automatically handles label conflicts and refreshes the display. This provides an interactive workflow to fix identity switches, merge broken tracks, or remove spurious detections directly within the software.

#### Segmentation model training

To train a model, users can select any combination of channels as the input. Each channel’s normalisation can be independently adjusted using either percentile or absolute normalisation. For RICM images, which are background-corrected with intensities centred around 1, an absolute normalisation range of 0.75 to 1.25 was adopted. In contrast, fluorescence images—where light source power and exposure often vary across experiments—use percentile normalisation, typically set between the 0.1-th and 99.99-th percentile for each image. During training, images in the training set are randomly augmented with operations such as flips and lateral shifts. Each input channel can undergo independent intensity adjustments, including blurring (random sigma between 0 and 4 px), physical noise addition (Gaussian, local variance, Poisson, and speckle), and random deactivation of channels (excluding the primary input channel), encouraging the model to learn redundancy across channels. StarDist models are trained for around 300 epochs with a learning rate of approximately 10*^−^*^4^ and a typical batch size of 8, while Cellpose models are trained over roughly 3000 epochs with a learning rate near 10*^−^*^2^ and a similar batch size.

#### Annotations for segmentation

To improve the annotation experience in napari, we developed a plugin that addresses common annotation errors identified with our expert annotators. The plugin detects repeated label values by comparing cell bounding boxes with mask areas, interpreting significant differences as multiple cells sharing the same label. Noncontiguous objects are automatically separated and relabeled, while small objects (less than 9 px^2^) are filtered out. The plugin also reassigns label values sequentially from 1 to the total number of objects, manages annotation naming and storage, and appends experiment metadata to each annotation, enabling precise tracking of annotation origin and context.

#### Classification and condition-based event detection

We developed a module to classify cells by writing conditions on the measurements with AND and OR logic gates. A flow-cytometry-like scatter plot showing the classification result at each time point for any combination of measurements assists the user in writing arbitrarily complicated conditions. This classification is instantaneous, as each cell from each time point is classified independently. With cell tracks available, we can perform a time-correlated analysis of the binary condition response for each cell. We implemented two particular descriptions: the first is that the cell has a unique state (condition always true or false): the median response determines the cell class; the second is the irreversible event, introduced to describe the cellular events in our applications. It implies three possible states for each cell: always false (no event observed), always true (the event happened before the beginning of observation), or transitioning from false to true (the event occurred during observation). For transitioning events, the software catches the event time by fitting a sigmoid over the binary response for each cell.

#### Deep learning event detection

To automate event detection, we implemented a deep learning approach with two convolutional models: (1) a classifier that assigns each cell to one of three event classes based on selected measurement time series, and (2) a regressor that estimates the event time for cells exhibiting the event, using the same time series input. Both models share a common backbone, a modified 1D ResNet, designed to process multivariate time series (Supplementary Fig. S7). To match the user-defined input tensor size, time series endpoints are extrapolated by repeating the edge values. In the training set, the time series are augmented with random noise and time shifts to balance the classes and artificially generate a quasi-uniform event time distribution. Users can opt to use our pre-trained event-detection models for specific applications (Tab. S4) or easily train new models on their own datasets generated within the associated viewer, with only a few clicks. Input time series are structured as tensors of shape (*T* × *n*_channels_), where *n*_channels_ represents the number of coupled time series (or channels) and *T* is an arbitrary maximum time series length, typically set to 128 frames. The input tensor is initially processed by a 1D convolution layer with a (1, ) kernel to expand it into 64 filters. It then passes through two 1D-ResBlocks, each with a (8, ) kernel and 64 filters, followed by a max-pooling layer (size 2) that doubles the number of filters to 128. The tensor continues through two additional 1D-ResBlocks before undergoing a global average pooling operation. A dense layer with 512 neurons collects the convolutional information, followed by a dropout layer with a 10 % dropout rate. The final layers differ by model task: for classification, a layer with three neurons and softmax activation distinguishes between three classes; by convention “event observed”, “no event observed”, and “event left censored”; for regression, a single neuron with linear activation predicts *t*_event_. The models are trained for 300 and 600 epochs, respectively, using the Adam optimiser with a user-controlled learning rate (typically 10*^−^*^4^, unless mentioned otherwise). For classification, categorical cross-entropy loss is minimised, with class weights introduced to mitigate class imbalance. For regression, the mean squared error is minimised. The batch size, usually set to 64 or 128 depending on the dataset size, is user-controlled. Only cells classified as the first class (“event observed”) proceed to the regression model during both training and inference.

#### Neighbourhoods

Single cells from two populations can be linked through a neighbourhood scheme, which can either define relationships within a single cell population (e.g., targets-targets) or across two distinct populations (e.g., effectors-targets). To accommodate specific application requirements, we developed two distinct neighbourhood methods. The mask contact method identifies touching cell masks at any time point, using mask dilation to allow a tolerance for cell contact. The isotropic method, by contrast, disregards cell masks and instead uses a circular region of a selected radius, centred on a reference cell’s centroid, to determine neighbouring cells. This approach is particularly useful for quantifying cell co-presence in cases with nuclear segmentation or unreliable masks. Neighbour counts extend the single-cell data tables in the same way as other single-cell measurements (more details in Supplementary Fig. S8). The reference-neighbour pairs identified at least once are then measured (e.g., relative distance, angle, velocity) and stored as rows in a cell-pair table. Similar to single cells, these pairs can be classified and annotated through an interactive cell pair viewer, which will be demonstrated in the second application.

### Details on the analysis of the cell-surface assay

#### Image preprocessing

A RICM background for each well of the cell-surface assay was reconstructed using the model-free background estimate method in Celldetective (described in Supplementary Fig. S3B). The reconstructed backgrounds were optimised and used to divide the RICM images, yielding float intensities centred around 1.

#### Single-cell extraction

The traditional segmentation pipeline consisted of two mathematical operations on the pixels (subtracting one and taking the absolute value) and a Gaussian filter (kernel of 0.8 pixels), followed by thresholding, watershed and automatic removal of non-lymphocyte objects. The pipeline was defined with the dedicated Celldetective module (Supplementary Fig. S4). To track the cells, the bTrack configuration was tuned to increase the isotropic spatial bin size to consider hypotheses (dist_thresh = 99.99), so that hypotheses could be generated for each tracklet. The branching, apoptosis and merging tracklet reconnection hypotheses were disabled and the segmentation miss rate reduced to 1 %. The position-based intensity measurements were performed in a circle of 2 *µ*m diameter centred around the cell positions.

#### Spreading rate

For each spread cell, the area was differentiated as a function of time using a sliding window of 5 frames (1 min 18 s) looking forward in time: from *t_i_*to *t_i_*_+4_ (this was configured graphically in the table exploration feature of Celldetective). The value at the onset of spreading *t*_spread_ was then extracted for each cell (with a track collapse option in the software) and plotted.

#### Segmentation benchmark

To use the Cellpose *cyto3* model, we passed a diameter value of 15 *µ*m, a cellprob_threshold of 0 and flow_threshold of 0.4. For each sample in the test set, the lymphocyte detection accuracy was defined as: *DA* = TP*/*(TP + FP + FN), with TP the number of lymphocytes predicted, FP the number of non-lymphocyte objects detected and FN the number of lymphocytes missed. A detection was accepted when the intersection over union (IoU) reached at least 0.1. The segmentation quality was defined as the IoU between the ground truth and predicted masks when they could be matched (TP detections). Since hovering cells had an ill-defined area, the segmentation quality was only computed for the subset of spread cells. The scores reported in Fig. 2 are the average over all test samples of the lymphocyte detection accuracy and the segmentation quality. The average score was defined as the average of these two scores for each segmentation method. The benchmark was performed in a Jupyter Notebook.

### Details on the analysis of the cell-cell assay

#### Image preprocessing

The fluorescent bsAb and LAMP1 channels were corrected for the background using the paraboloid fit technique described in Supplementary Fig. S3A. The fitted background for each image was subtracted from the original image, without clipping. The movies from the LAMP1 experiment were aligned prior to analysis with the SIFT multichannel registration plugin in Fiji (to be able to investigate effector cell velocities). The macro used to align the movies in a Celldetective experiment project is available in the documentation.

#### Tracking of target cells

The MCF-7 nuclei were tracked with a quasi-stationary motion model, allowing for small displacements and potentially long time gaps. We passed the Hoechst and PI intensities and the nuclear apparent area to the tracker, which helped in experiments where the live/dead transitions were not too sudden. Cells not present in the first frame of the movie were filtered out to limit the number of false positive detections and exclude daughter cells from our study. Tracks were extrapolated until the end of the observation to measure intensities at the last detected position, which worked well in this system since the nuclei were mostly stationary.

#### Effector cell extraction

For the dynamical study, manual corrections of segmentation masks were performed in napari (see Supplementary Fig. S5) for both target and effector populations. These corrections included splitting merged masks of adjacent cells, removing false-positive detections (e.g., debris or out-of-focus cells), and adding missed cells. This was necessary to achieve tracking quality sufficient for reliable pair analysis. Effector cell velocity was computed over a sliding window of 3 frames (5.19 min) centred on the current time (from *t_i−_*_1_ to *t_i_*_+1_).

#### bsAb fluorescence measurements

The mean bsAb fluorescence intensity over each NK cell mask was measured from the background-corrected images.

#### LAMP1 measurement

The mean LAMP1 intensity over each NK cell mask was measured from the background-corrected anti-LAMP1 APC fluorescence images. A representative image of LAMP1 antibody staining is shown in Supplementary Fig. S10. We classified cells as LAMP1-positive or LAMP1-negative at each time point using an intensity threshold (*I*_LAMP1_ *>* 5 a.u.).

#### Contact classification

Target cell contact with NK cells was inferred using a radial neighbourhood calculation, as only target cell nuclei masks were available, and was further classified by the number of neighbouring target cells: an NK cell in contact was defined as having at least one target neighbour (inclusive counting convention, see Supplementary Fig. S8). A quality check of this contact classification was performed in-situ on the images using a dedicated viewer that displays cells directly on microscopy images (Video single-timepoint-contact.mp4).

#### Simulation

The Monte-Carlo simulation for the sum of bsAb fluorescence signals and its comparison with the other conditions was performed in a Jupyter Notebook with in-house code and functions from the Celldetective package. This notebook allowed us to produce Figure 4C and Supplementary Fig. S9.

#### Segmentation benchmark

Nuclear detection accuracy was defined in the same way as the lymphocyte detection accuracy in the cell-surface assay. To use the Cellpose *nuclei* model, we set the diameter to 22 *µ*m, the cellprob_threshold parameter to 0.0 and the flow_threshold parameter to 0.4. The segmentation quality is computed over all nuclei. As before, the scores reported in Fig. 3 are the average over all test samples of the detection accuracy and segmentation quality. The average score was defined as the average of these two scores for each segmentation method.

#### Death event detection benchmark

The event detection models we present were trained on the db-si-NucPI dataset available in Zenodo with a 20 % validation split and 10 % test split, randomised at each training. For each tested combination of batch size (BS ∈ [64, 128, 256]) and learning rate (*α* ∈ [10*^−^*^4^, 10*^−^*^3^, 10*^−^*^2^, 10*^−^*^1^]), we trained 10 event detection models for 300 epochs with the two different input approaches (PI time series alone or PI + nuclear area time series). The time series were normalised with a percentile normalisation (0.1 % and 99.9 %) and clipping. The model input tensor length was set to 128 frames. For the hyperparameters that achieved the best average performance (here BS = 64 and *α* = 10*^−^*^4^), we picked the best event detection model produced out of the 10 for the benchmark of Fig 3D. This benchmark was performed on a separate test set, identical for all methods. The annotations came from 3 movies, two of which belonged to experiments never seen by the models. The third is from a position adjacent to a position seen by the models. It was selected due to its over-representation of cells that are already dead from the beginning, or dying at the very beginning, which are very rare in most experiments. The benchmark was performed in a Jupyter Notebook.

### Statistics

#### Statistical tests

Since for all multi-condition comparisons at least one distribution was not normally distributed, we assessed statistical difference using the nonparametric two-sample Kolmogorov-Smirnov one-sided test (Python package scipy.stats.ks_2samp). P-values were converted into star notations following the GraphPad convention: P *>* 0.05: ns, P *<* 0.05: *, P *<* 0.01: **, P *<* 0.001: ***, P *<* 0.0001: ****.

#### Effect sizes

For statistically significant differences, we computed Cliff’s Delta to report the effect size (Python package cliffs-delta). We followed the conventions of the package to convert the effect size value into a qualitative effect: *δ <* 0.147: negligible, *δ <* 0.33: small, *δ <* 0.474: medium, *δ >* 0.474: large.

#### Plots

Most plots presented in this article were generated within Celldetective using the matplotlib and seaborn packages. Notable exceptions include the segmentation benchmark barplots, confusion matrices, and correlation plots (Figs. 2B,E and 3B,D), which were produced in Jupyter notebooks using matplotlib and scikit-learn. Composition was done in Inkscape to add statistical tests and effect sizes or merge separate plot layers.

## Acknowledgements

We thank Cristina Gonzalez and Maddy Messa N’Dong for preliminary experiments; Juliette Prothon for producing and labelling the bispecific antibodies; Adrien Aimard, Dominique Touchard, Martine Biarnes for excellent technical support; Gaurav Verma for multiple bug reporting; Pierre-Henri Puech and Thierry Galliano for fruitful discussions. The project leading to this publication has received funding from France 2030, the French Government programme managed by the French National Research Agency (ANR-16-CONV-0001) and from the Excellence Initiative of Aix-Marseille University - A*MIDEX AMX-21-IET-017, as well as AMIDEX Emergence Innovation (project ForSelecAntibody), Plan Cancer PhysCancer programme (project ComPhysAb) and ANR-22-CE09-0014.

## Supplementary Figures

**Video: ricm events.mp4**

**Video: adcc rgb.mp4**

**Video: single-timepoint-contact.mp4**

**Video: cell interactions.mp4**

---

**Fig. S1.**
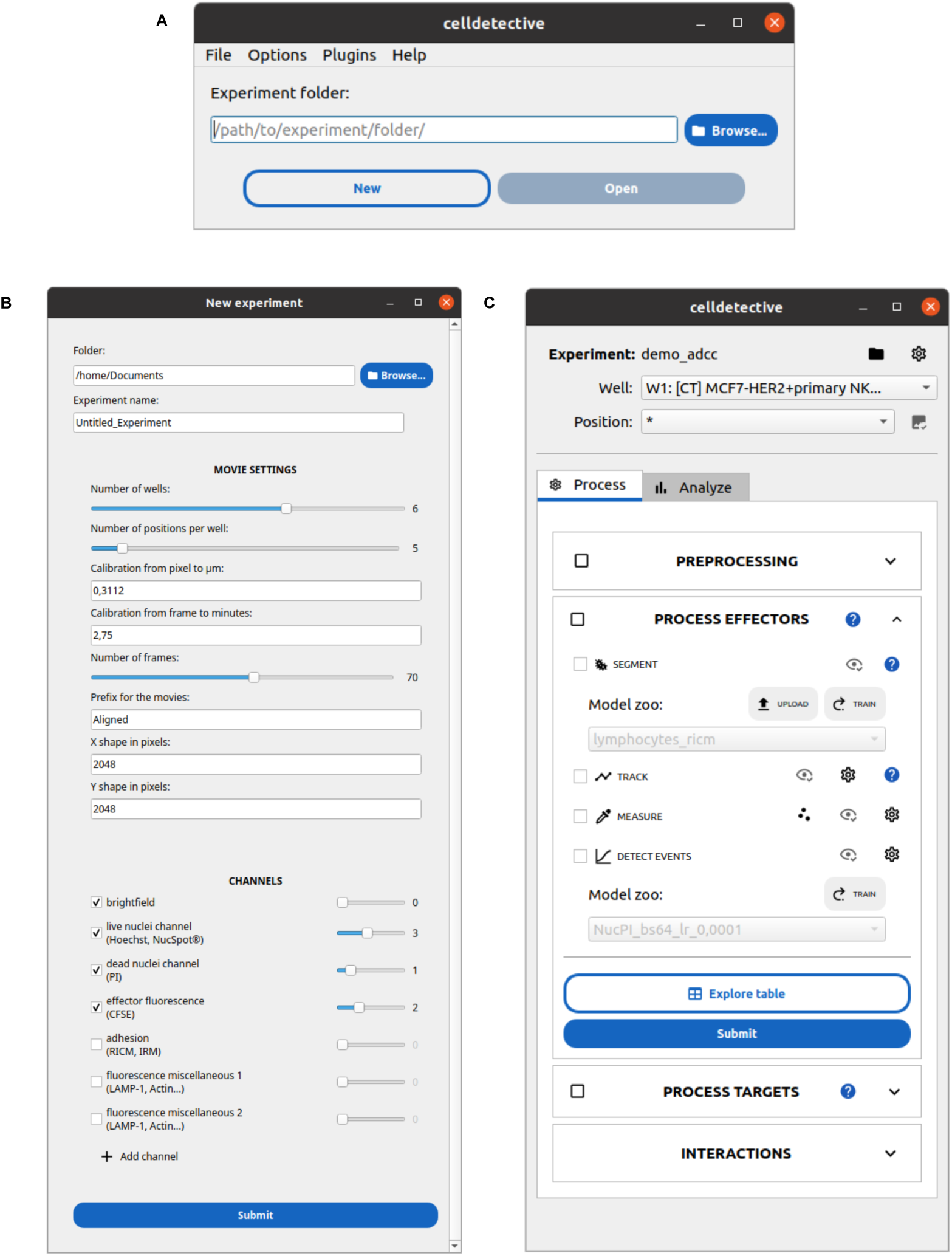
Main GUI windows: A) Experiment project selection window. B) GUI to generate a new experiment, where the user provides the metadata. C) Main interface after loading a project. The process block for the effector population is unraveled, showing the 4 main steps detailed in Fig. 1.

**Fig. S2.**
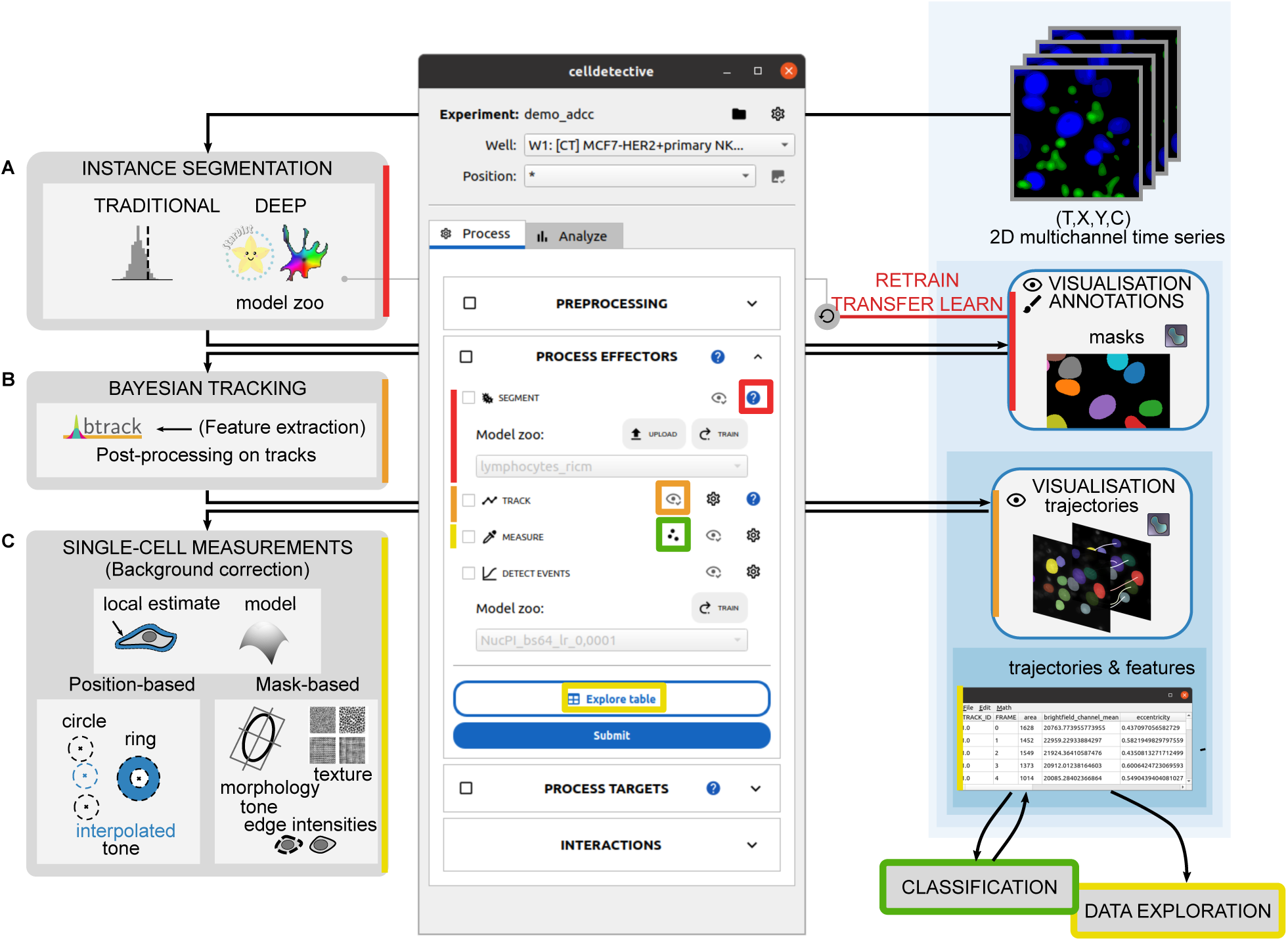
Processing modules to extract single cells: Processing modules are shown on the left, with brief illustrations of the main options, while the output and visualisation modules are represented on the right side, following Celldetective’s graphical structure. A typical pipeline features successively (from top to bottom): A) an instance segmentation (either with traditional thresholding method or a deep learning model, stored in a model zoo, using StarDist [36] or Cellpose [38]). The output data are masks, which can be visualised and annotated using napari [57], for correction or model retraining. B) Bayesian tracking associated with feature extraction (btrack [42]), the output data being trajectories, which can also be visualised with napari. C) Single-cell measurements of intensity, morphology, texture (based on cell position or on mask), the output being tables of trajectories enriched with features.

**Fig. S3.**
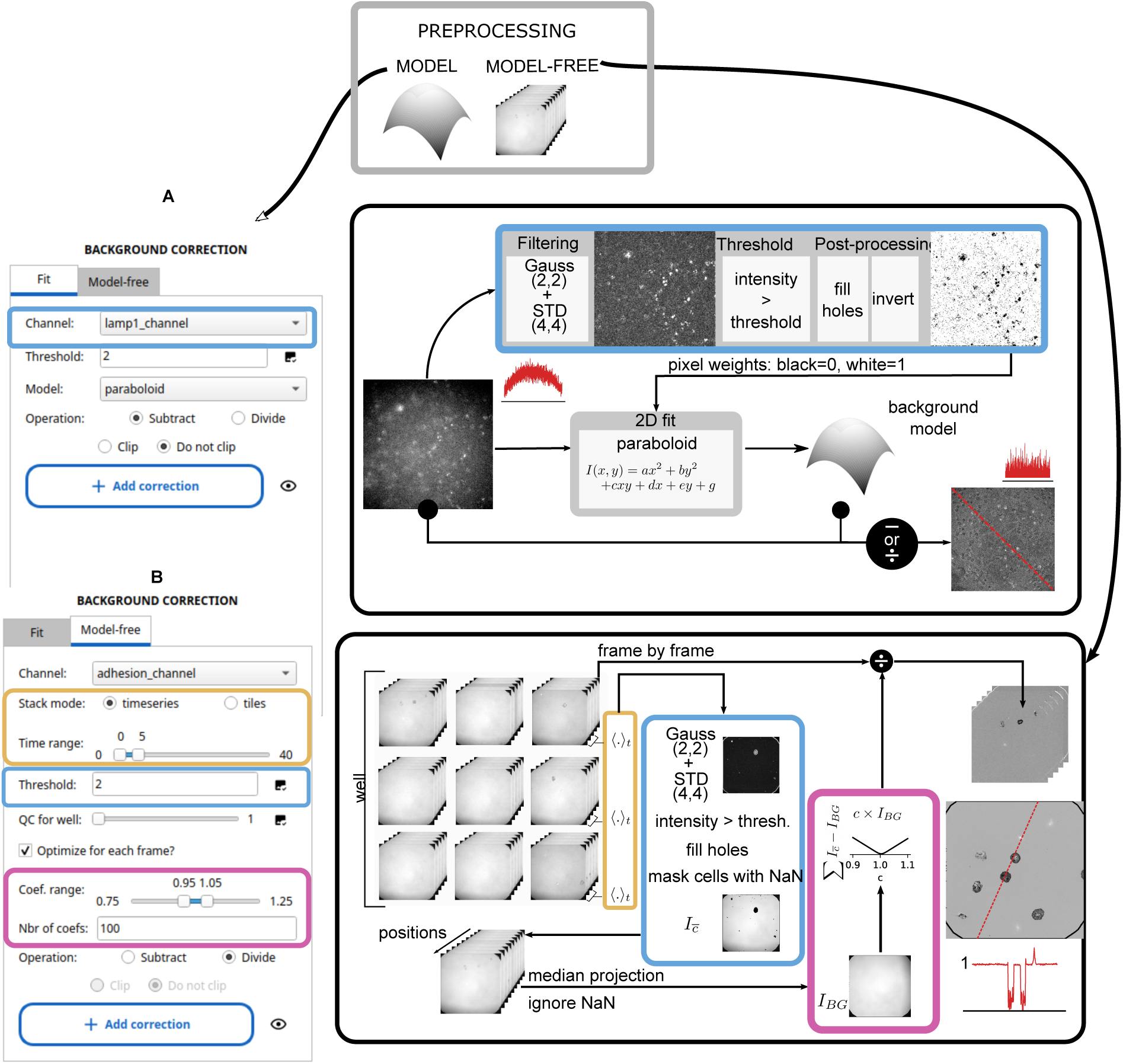
Background correction methods. Screenshots of the background correction tabs in the software and illustration of the associated pipeline. Coloured bounding boxes match graphical parameters to their role in the pipeline. The thick black arrows indicate the starting point of the pipelines. A) The steps for a fit-based correction involve selecting a channel of interest, setting a threshold on the standard-deviation filtered image to exclude the cells (done graphically with a viewer). Then a model (either plane or paraboloid) is fit on the background pixels for each image. The fitted background is either subtracted or divided, with or without clipping. B) With the model-free approach, a single background is reconstructed per well, following the same threshold approach but leveraging the multi-positional information to construct a median background. Instead of applying the background directly, it is amplified to minimise the difference with the current image’s background. As with A), the background is then either subtracted or divided, with or without clipping.

**Fig. S4.**
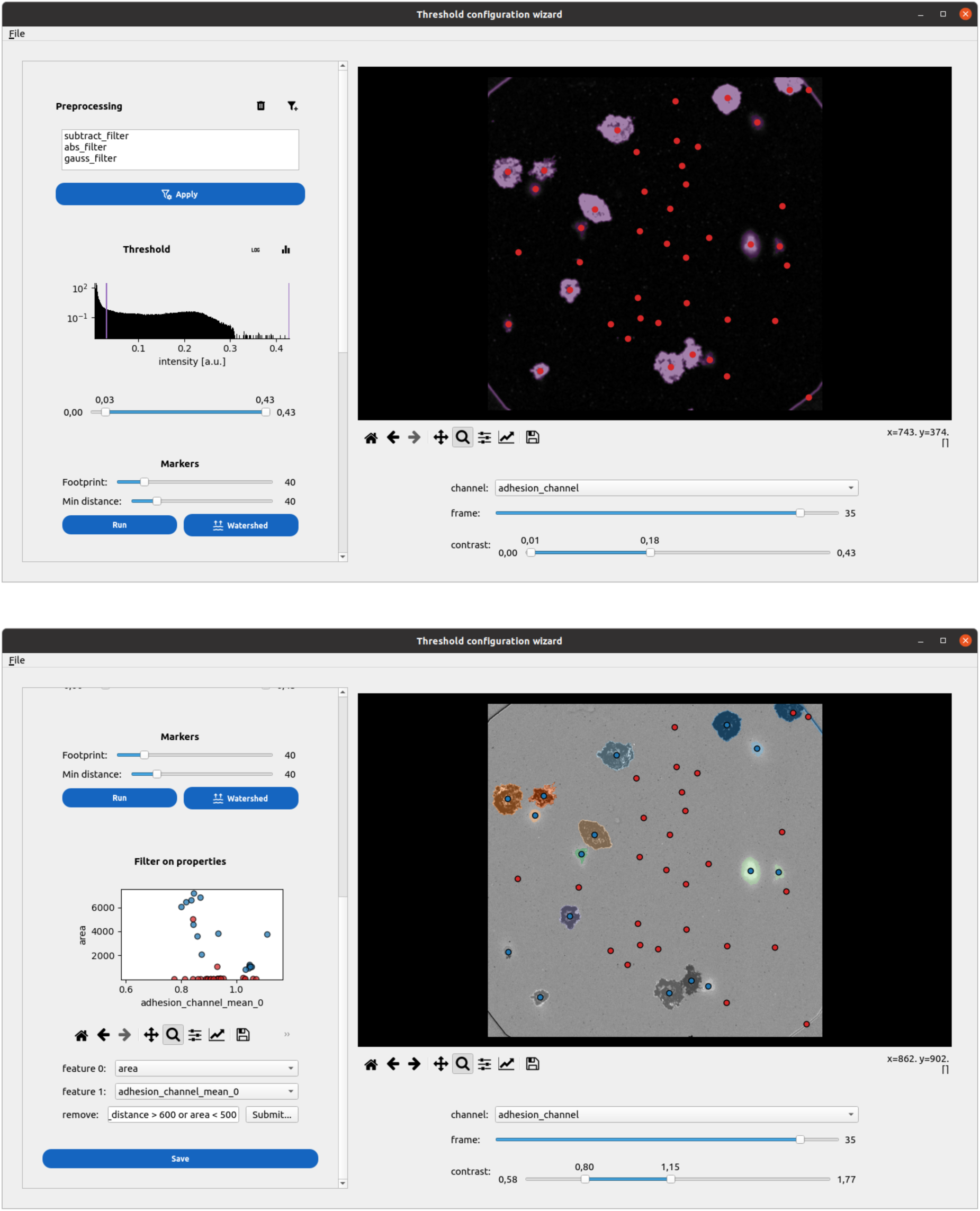
Designing a traditional segmentation pipeline. Screenshots of the traditional segmentation pipeline configuration interface applied to a normalised RICM image of spreading primary NK cells. Top: User-selected mathematical operations (subtraction, absolute value) and preprocessing filters (Gaussian blur) are applied to the image, with results displayed in real-time on the right. A binary mask, set using an interactive thresholding slider, is overlaid on the transformed image in purple. Spot detection is then performed on the Euclidean Distance Transform of the binary mask, using parameters for footprint size and minimum object distance; detected spots are shown as red scatter points. Bottom: After applying the watershed algorithm, the instance segmentation of cells is overlaid on the original normalised RICM image. The user can set filters on object features to exclude false positives; rejected cells are marked in red in both the feature scatter plot and the image.

**Fig. S5.**
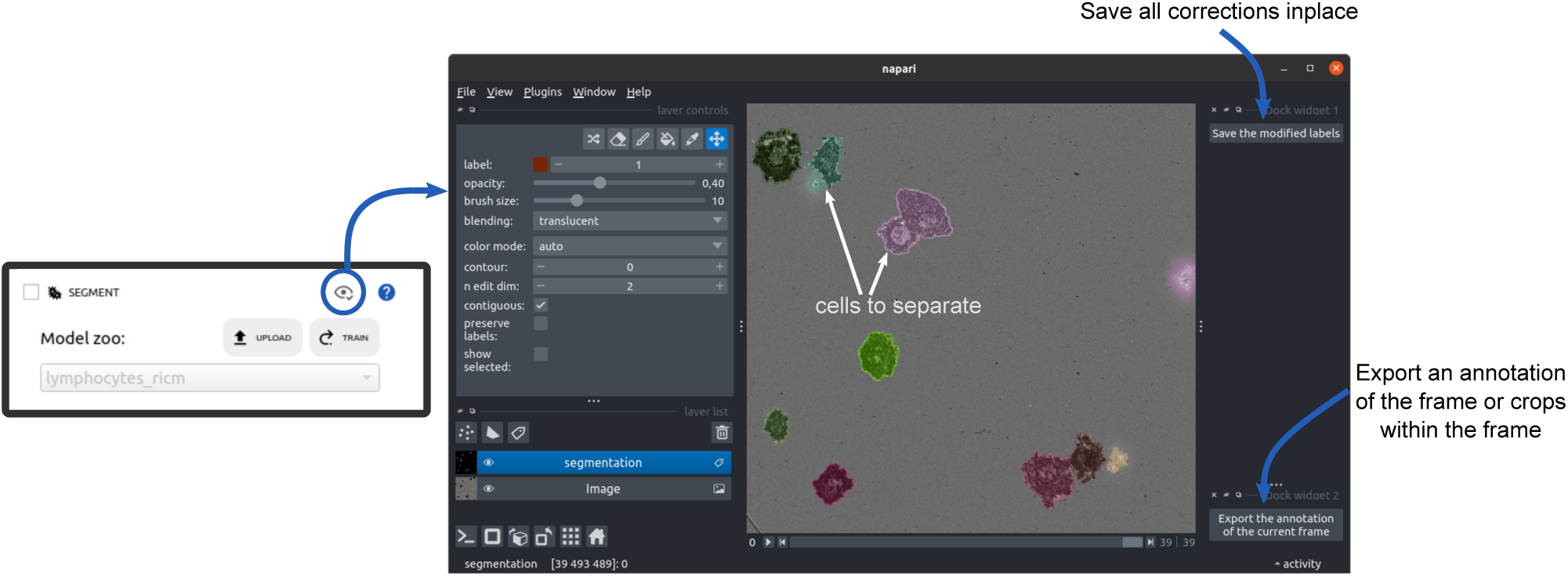
Segmentation correction and annotation. To view segmentation results, the user can click the “eye” button in the segmentation module, which opens images and labels in napari. Shown is a screenshot of a segmentation outcome for a normalised RICM image of spreading primary NK cells, with labels overlaid on the image. The user can correct the masks directly on the image, for example, to separate cells indicated by white arrows. New labels can be assigned and saved, and the full set of corrected masks can either be saved in place (top-right plugin) or exported as a training sample for a segmentation model (bottom-right plugin).

**Fig. S6.**
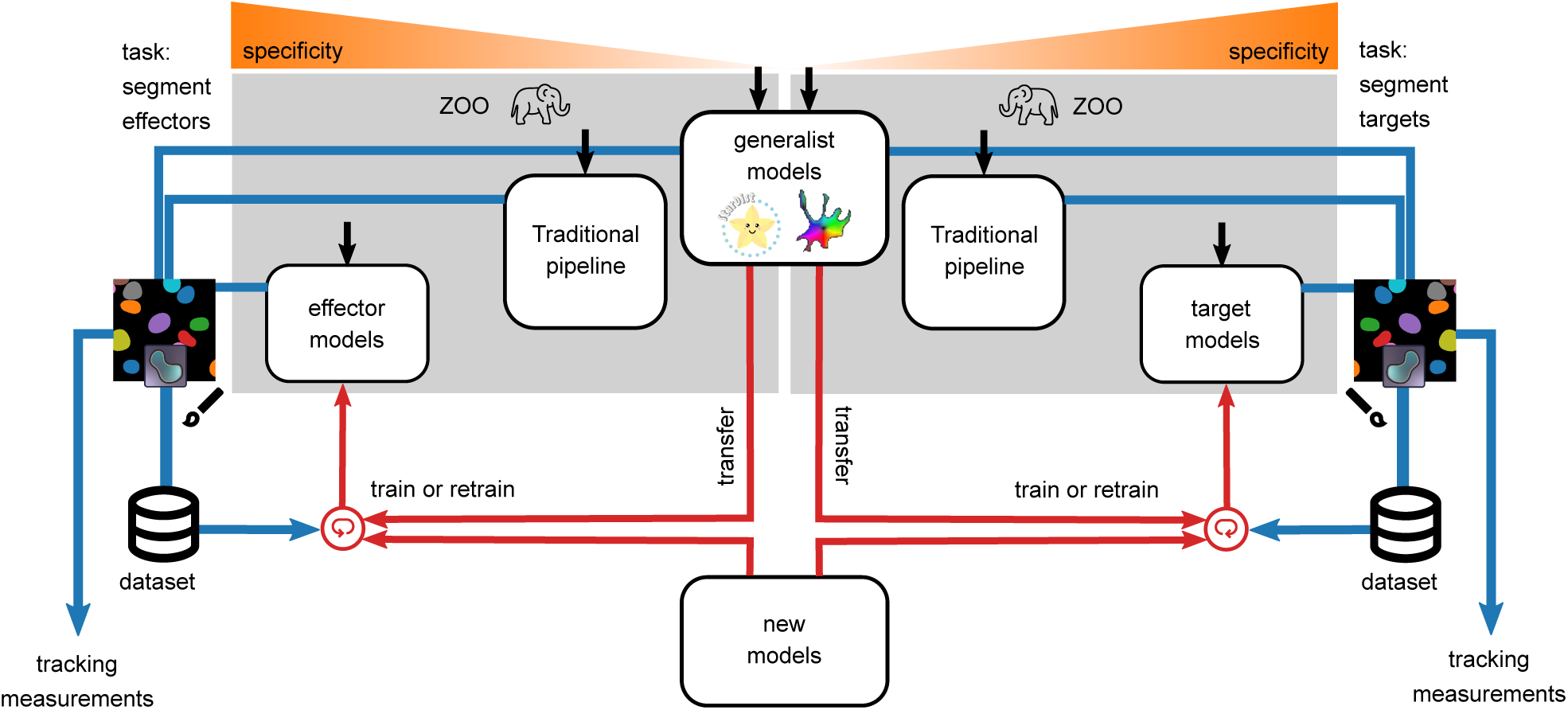
Overview of segmentation strategies. The principle is illustrated for a mixture of two cell populations, effector and targets. Celldetective provides several entry points (black arrows) to perform segmentation, with the intent of specifically segmenting a cell population (left: effectors, right: targets). The generalist DL models are listed in Tab. S1. Specific DL models are listed in Tab. S2 for targets and Tab. S3 for effectors. The traditional pipeline refers to a thresholding method accessible through the GUI (see Fig. S4) and which can serve as a starting point to prepare a dataset for training a new DL model. The masks output from each segmentation technique can be visualised and manually corrected in napari. Exporting those corrections into a dataset of paired image/masks can be used either to fit a generalist model (transfer learning) or train a new model from scratch.

**Fig. S7.**
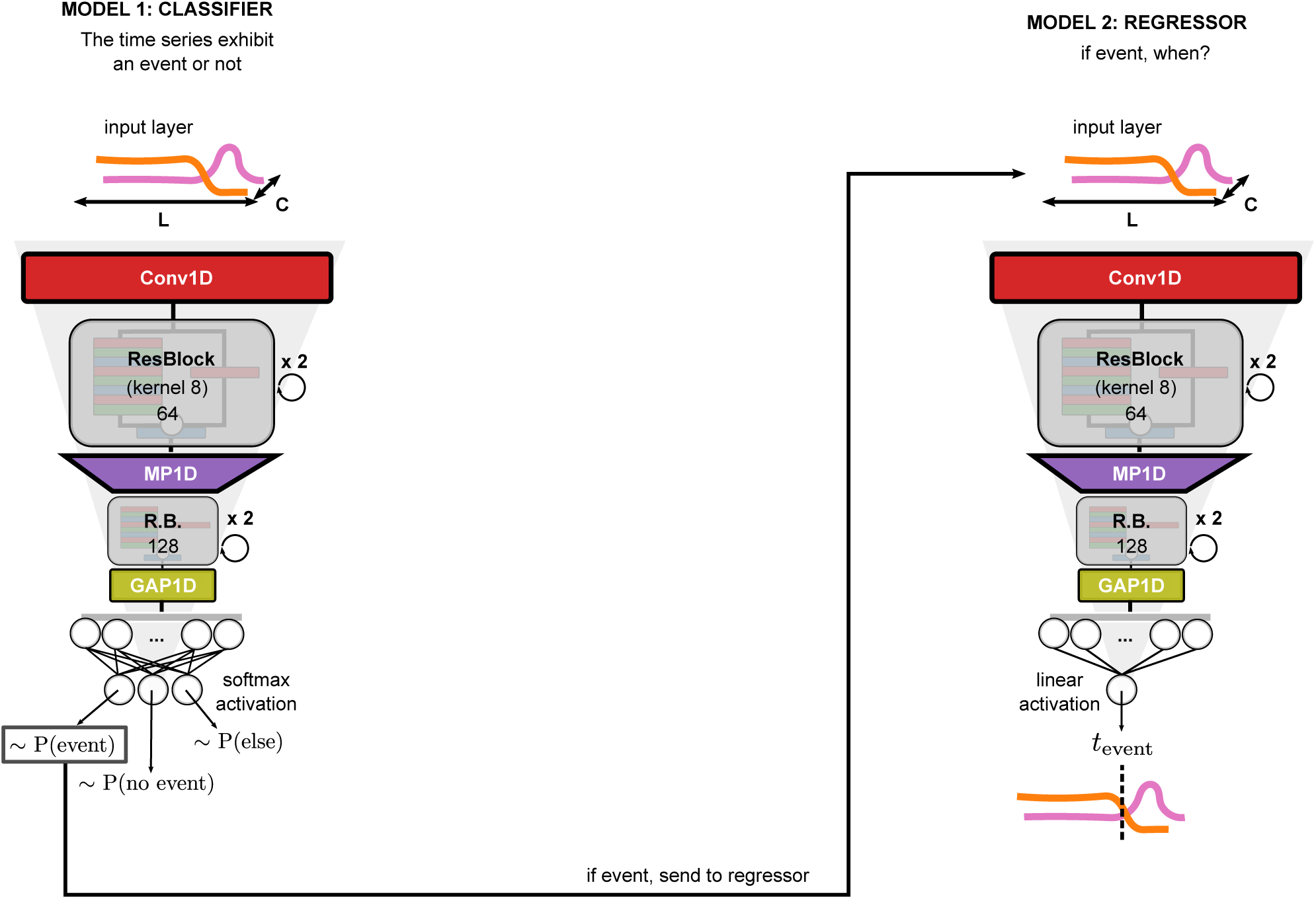
Event detection model architectures. The event detection models consist of two CNN-based models called back to back during inference to 1) classify single-cell time series and 2) detect the event times from the same time series. The models share a similar architecture of a first 1D convolution, followed by ResBlocks encoding the time series into either three neurons with a softmax activation for the classification problem (three classes) or one neuron with a linear activation for the regression problem. As illustrated here, only the time series that were classified as “event” are sent to the regressor model.

**Fig. S8.**
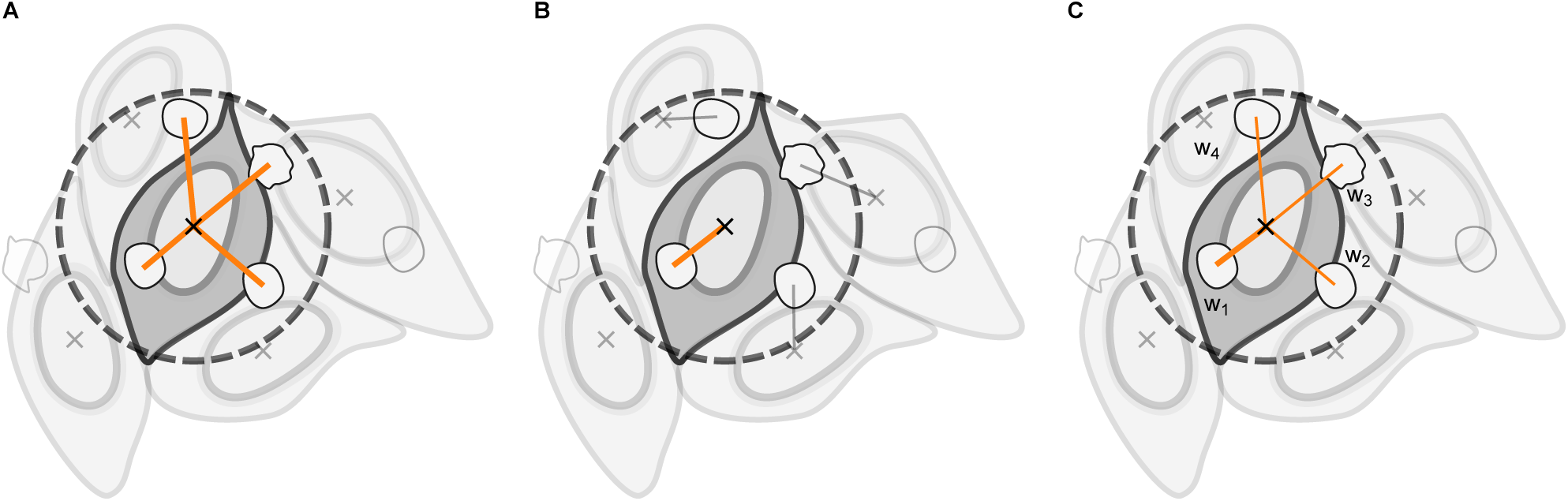
Neighbour counting methods: A) All neighbouring cells in contact or within the isotropic neighbourhood are linked to the reference cell (inclusive method). B) Each neighbouring cell is linked exclusively to the closest reference cell (exclusive method). C) Neighbouring cells in contact or within the isotropic neighbourhood are linked to the reference cell, with weights inversely proportional to their number of neighbours.

**Fig. S9.**
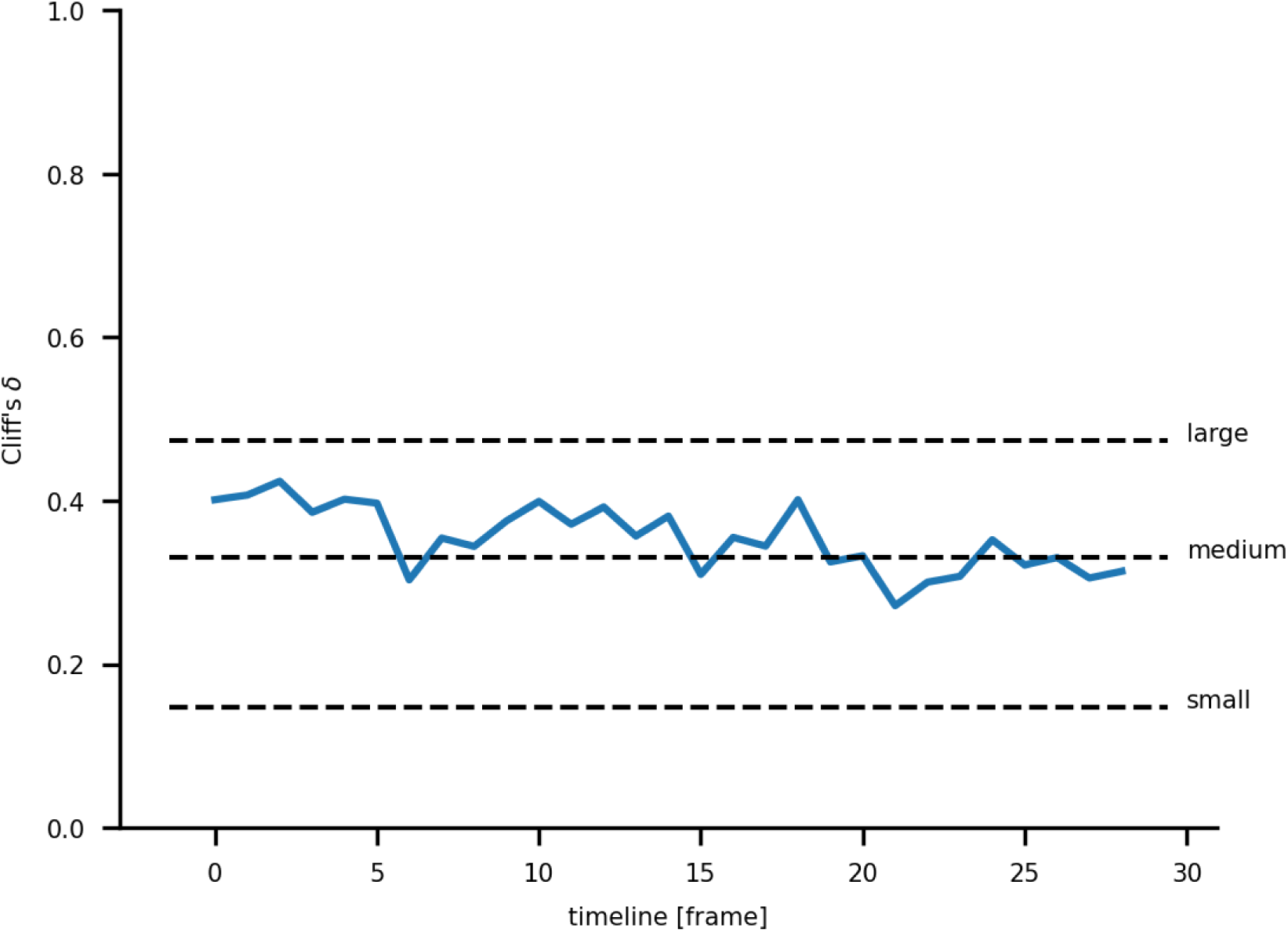
Effect size at each time point: Cliff’s Delta between the simulated sum of two distributions (CE4-21/off-contact/HER2+ + CE4-X/in-contact/HER2+) and CE4-21/in-contact/HER2+ computed at each timepoint, independently.

**Fig. S10.**
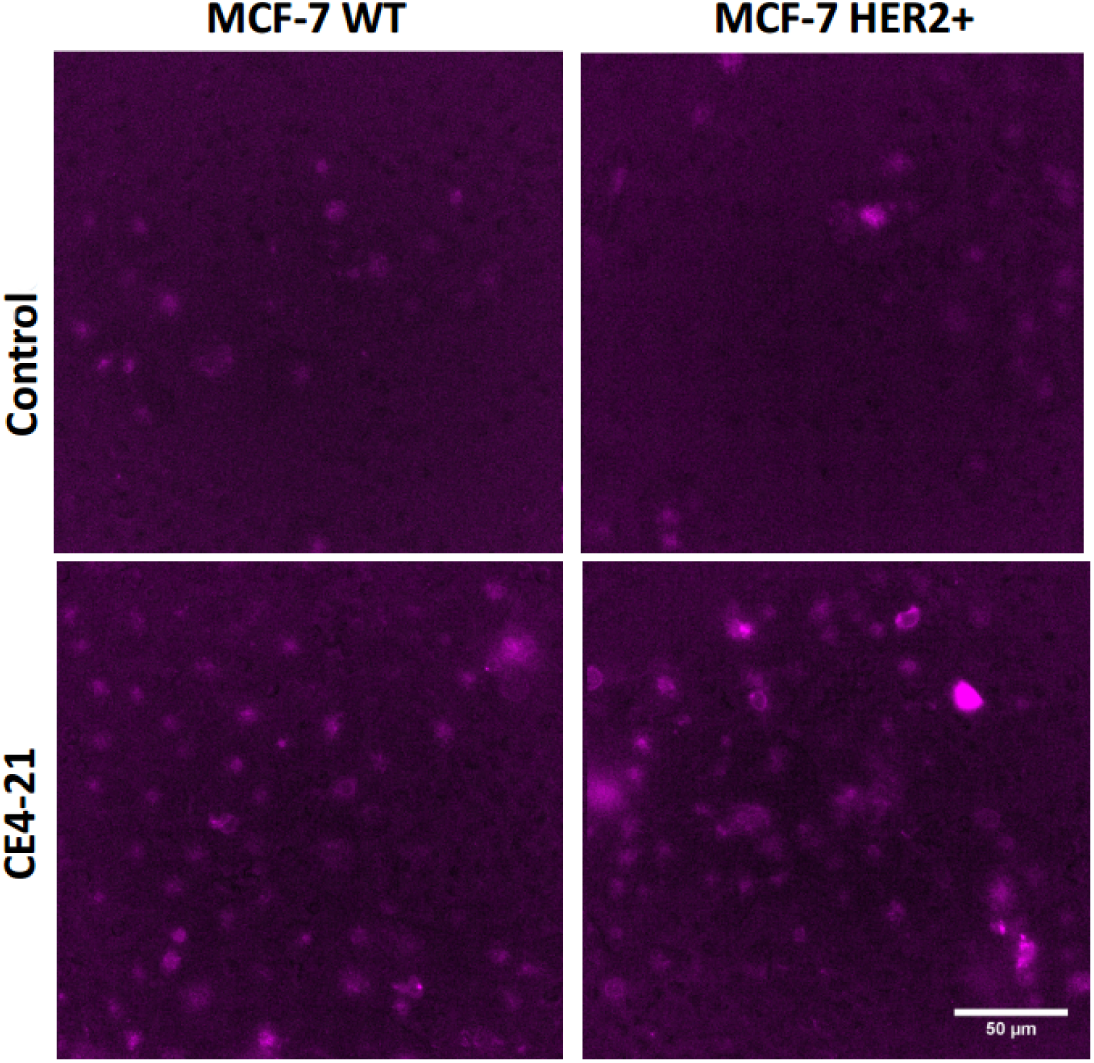
Representative LAMP1 antibody staining. Fluorescence image of the co-culture assay showing NK effector cells stained with anti-LAMP1 APC antibody. Top: Control without bsAb. Bottom: in the presence of 1nM bsAb CE4-21. Left: MCF7 WT targets. Right: MCF7-HER2+ targets.

**Table S1.**
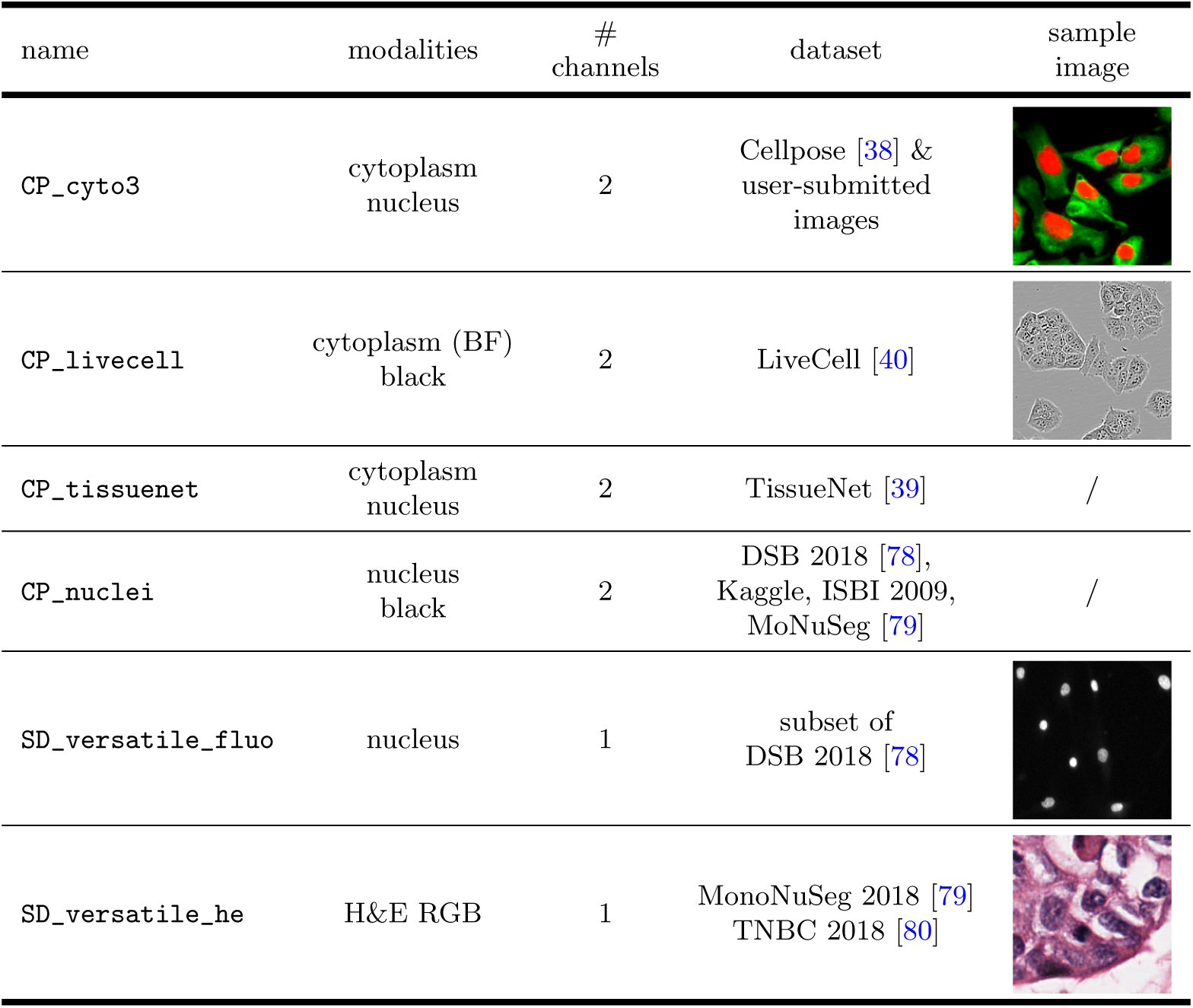
Generalist deep learning segmentation models. This table lists the different generalist models (Cellpose or StarDist) which can be called natively in Celldetective. The sample images are cropped to (200 *×* 200) px and rescaled homogeneously to fit in the table.

| name | modalities | #<br>channels | dataset | sample<br>image |
| --- | --- | --- | --- | --- |
| CP_cyto3 | cytoplasm<br>nucleus | 2 | Cellpose [38] &<br>user-submitted<br>images |  |
| CP_livecell | cytoplasm (BF)<br>black | 2 | LiveCell [40] |  |
| CP_tissuenet | cytoplasm<br>nucleus | 2 | TissueNet [39] | / |
| CP_nuclei | nucleus<br>black | 2 | DSB 2018 [78],<br>Kaggle, ISBI 2009,<br>MoNuSeg [79] | / |
| SD_versatile_fluo | nucleus | 1 | subset of<br>DSB 2018 [78] |  |
| SD_versatile_he | H&E RGB | 1 | MonoNuSeg 2018 [79]<br>TNBC 2018 [80] |  |

**Table S2.**
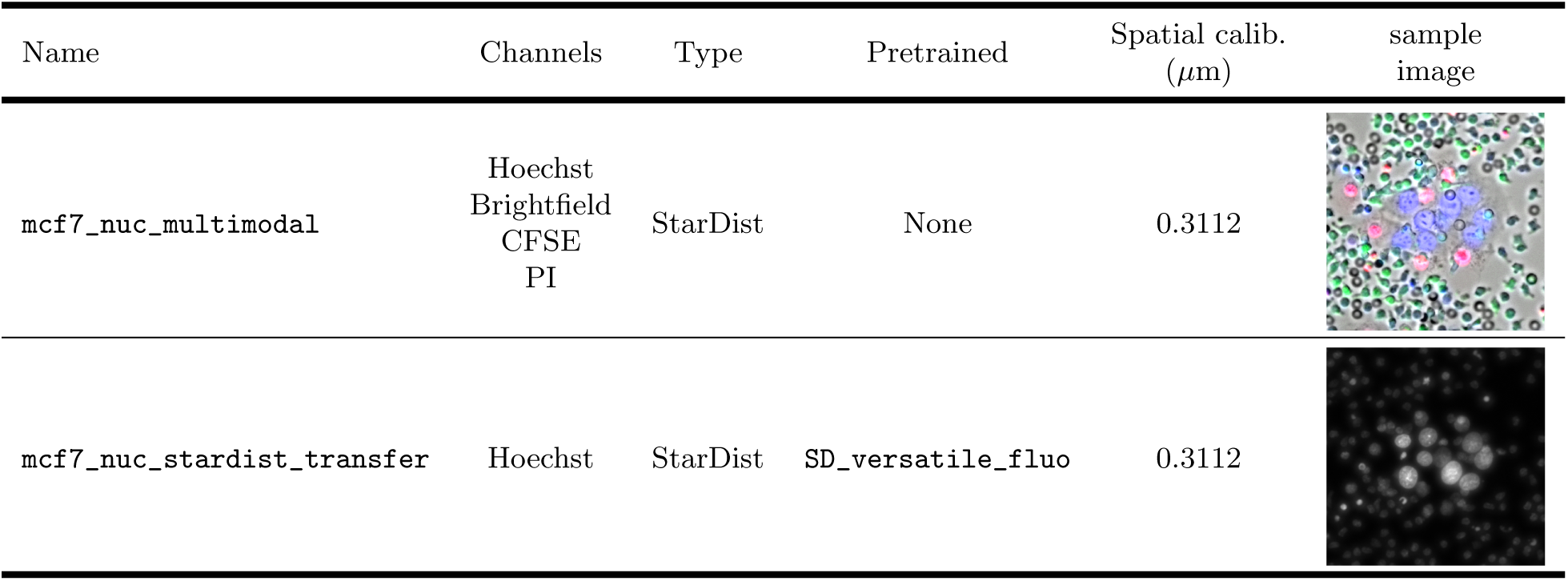
MCF7 nuclei segmentation models in the presence of primary NK cells. Each model was trained on the same dataset of ADCC images, picking only the relevant channels.

| Name | Channels | Type | Pretrained | Spatial calib.<br>($\mu\text{m}$) | sample<br>image |
| --- | --- | --- | --- | --- | --- |
| mcf7_nuc_multimodal | Hoechst<br>Brightfield<br>CFSE<br>PI | StarDist | None | 0.3112 |  |
| mcf7_nuc_stardist_transfer | Hoechst | StarDist | SD_versatile_fluo | 0.3112 |  |

**Table S3.**
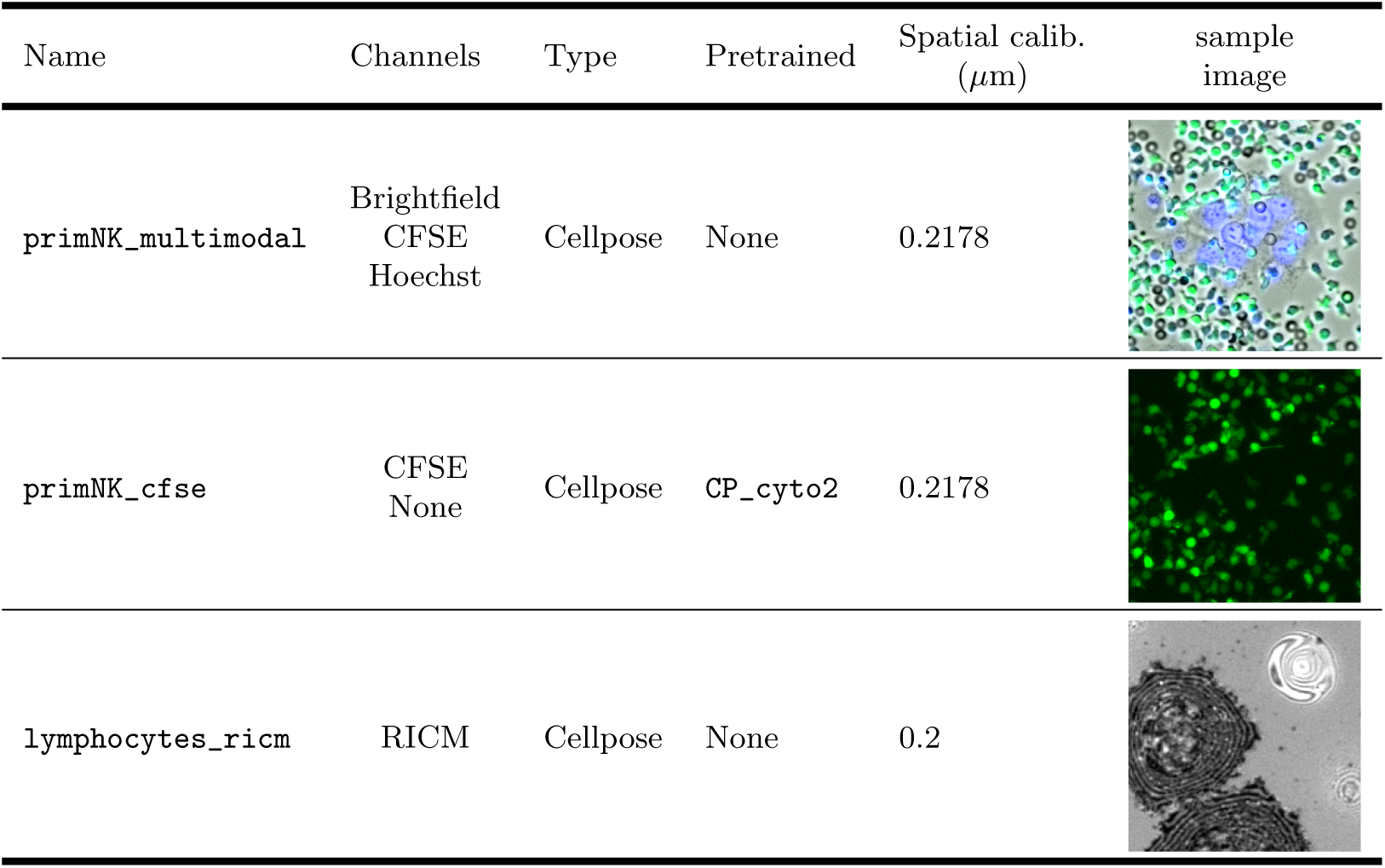
Primary NK segmentation models. The models have been trained on a dataset of annotated primary NKs in ADCC images (*primary NKs w MCF7*).

| Name | Channels | Type | Pretrained | Spatial calib.<br>($\mu\text{m}$) | sample<br>image |
| --- | --- | --- | --- | --- | --- |
| primNK_multimodal | Brightfield<br>CFSE<br>Hoechst | Cellpose | None | 0.2178 |  |
| primNK_cfse | CFSE<br>None | Cellpose | CP_cyto2 | 0.2178 |  |
| lymphocytes_ricm | RICM | Cellpose | None | 0.2 |  |

**Table S4.**
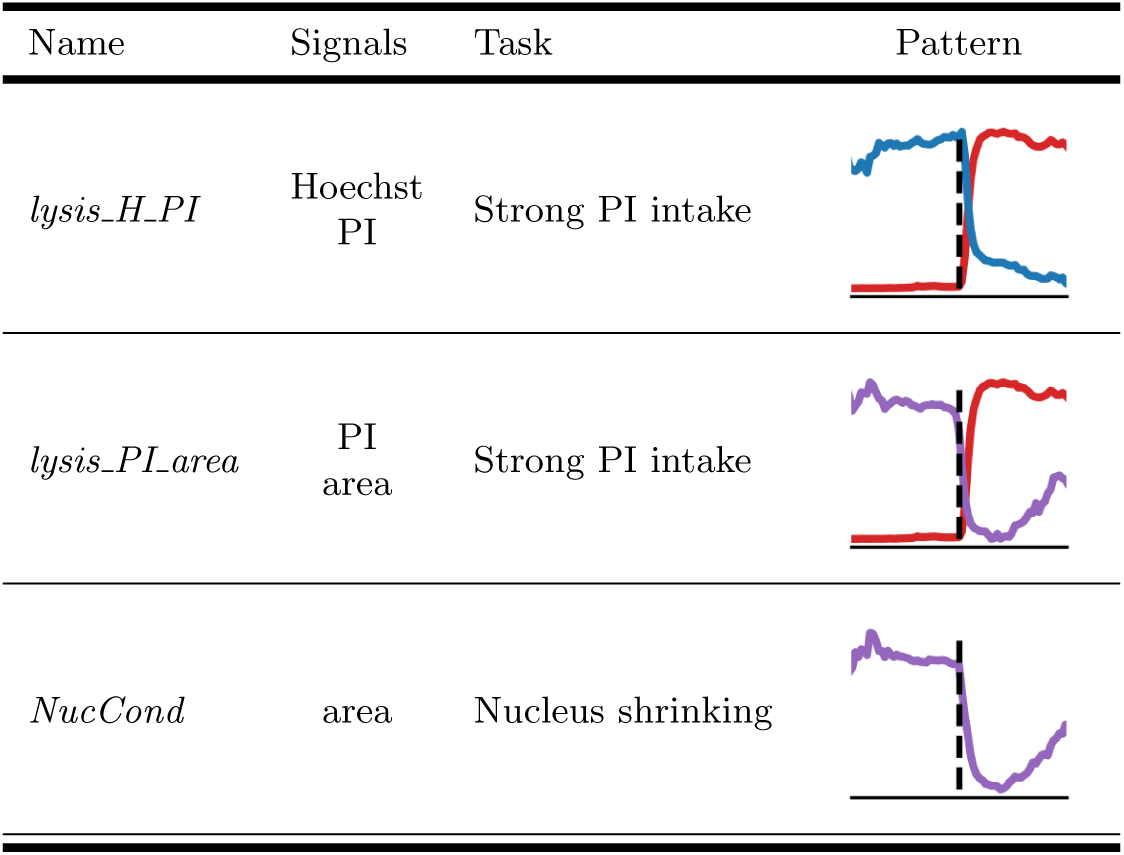
Event detection models. We trained the following 1D DL models to classify and regress events of interest. The mean event response, centred at the event time, is shown for each channel in the pattern column.

